# Modeling phage therapy

**DOI:** 10.1101/2025.11.08.687339

**Authors:** Rob J. de Boer, Robert Schooley, Alan S. Perelson

## Abstract

Patients infected with life-threatening multi-drug resistant (MDR) bacteria have been treated with cocktails of bacteriophages. This is a complicated form of personalized medicine as the phages given to a patient have to be selected beforehand on the basis of their lytic capacity of the infecting bacteria. Because bacteria rapidly become resistant, the evolution of resistance to a diverse cocktail of phages is a complicated dynamical process, during which competing bacterial strains replace one another by accumulating several resistance mechanisms, each of which may involve a fitness cost. As a consequence, it is typically not known why a particular phage therapy succeeded or failed, and how one can optimize the composition of the cocktails to maximize the rate of success.

To improve upon this, we extend an existing *in vivo*-calibrated mouse model into a novel mathematical model for the human situation, and include multiple phages infecting multiple bacterial strains, differing in their resistance to each of the phages. We adjust several parameter estimates of the bacterial model to the human situation, and use the model to describe a successful case of phage therapy involving several cocktails, each containing several phages. In the model, treatment success crucially depended on pretreatment resistance levels, and on the diversity and the timing of the cocktails. Once an appropriate cocktail is found, it is less important to further optimize the infection rates of the phages. Resistant bacterial strains expand rapidly when sensitive strains decline, and the higher the infectivity of the phages, the faster resistant strains expand. Because resistance evolves rapidly, it is best to provide a diverse set of phages right from the start of therapy, i.e., to hit hard and early, and create a high genetic barrier to bacterial resistance.

## 1 Introduction

Due to the evolution of bacterial resistance to multiple types of antibiotics, viruses that can infect and kill bacteria (called bacteriophages), are receiving renewed interest as a treatment option for bacterial infections. Recent clinical successes with patients recovering from rampant bacterial infections due to personalized phage therapy, have sparked interest into the design, efficacy, and safety of cocktails of phages that can be given to patients [12, 24]. Phage therapy has been used as ‘living antibiotics’ in the Soviet Union over extended periods of time [29], but these recent successes were ‘compassionate use’ cases of life threatening bacterial infections that were treated with personalized cocktails of phages hastily selected based on their capacity to lyse the bacteria causing the infection [6, 8, 20, 28]. Unfortunately, the very limited host range of most bacteriophages calls for such a personalized approach, and is a major challenge for generalized treatments and clinical trials [12].

Phage therapy may work for several reasons: (1) phages may simply infect and eliminate all bacteria (which is unlikely due to the rapid evolution of resistance [5, 16]), (2) phages may reduce bacterial concentrations to levels where the host immune system regains control [26], (3) by evolving resistance to phages bacteria may regain sensitivity to particular antibiotics [6, 24, 27, 28], (4) by modifying their capsule to prevent infection [20] bacteria may become more sensitive to killing by neutrophils, and (5) bacteria that evolved phage resistance may have such a crippled fitness that their replication rate drops below the rate at which the host immune system eliminates them. It seems virtually impossible to distinguish between these mechanisms on the basis of observational clinical data alone [8, 12]. Because we expect many more people with multiple drug resistant (MDR) infections to be treated with phage therapy in the near future [24], we here develop mathematical models implementing these mechanisms, with the aim to employ these models for better interpretation of future data. Such models may help one optimize the design of therapy, and by fitting variations of these models to the data, one may become able to identify the mechanism(s) underlying the control of rampant bacterial infections by phage therapy in individual cases.

The infection of bacteria by phages, and the rapid evolution of resistance by bacteria, has been studied extensively *in vitro*, and it is known that these systems tend to a approach an attractor where susceptible and resistant bacteria co-exist with the phages [5, 17]. It was therefore surprising that phage therapy can be successful *in vivo*, and by elegantly combining modeling with experiments in mice, Roach *et al*. showed that the innate immune response, in this case mostly neutrophils, is largely responsible for the *in vivo* success of phage therapy in their experiments [17, 26]. Apparently, neutrophils are usually incapable of controlling rampant bacterial infections, and a major effect of phage therapy is to reduce bacterial populations to a level where the killing of bacteria by neutrophils suffices to control bacterial growth. This mechanism requires the existence of a threshold in the bacterial density above which neutrophils can no longer control bacterial growth. Such a threshold is readily brought about in a model where at high bacterial densities (1) the neutrophils approach a maximum density, and (2) the number of bacteria killed per neutrophil per unit of time approaches a maximum, e.g., due to the fact that killing takes time [14, 23, 26].

Neutrophil densities rapidly increase during inflammation [25]. Because they are released more rapidly from the bone marrow, they can approach a new (higher) steady state level on a time scale of less than a day [1, 33]. Thus, there should be two thresholds: (1) the maximum bacterial density that can be controlled by the normal density of neutrophils, e.g., small bacterial invasions that are controlled on a daily basis without triggering additional inflammation, and (2) the much higher threshold above which infections can grow uncontrolled despite the massive killing by neutrophils at their maximal density. In between an innate immune response is required to control the infection before it breaches the upper threshold.

After developing a novel mathematical model having both thresholds, and allowing for several bacterial strains differing in their sensitivity to various phages and antibiotics, we illustrate its usefulness by describing a case of successful phage therapy of a patient suffering from a rampant infection with multi-drug resistant (MDR) *Acinetobacter baumannii* bacteria [28, 29, 32]. We present a few scenarios suggesting why this particular phage therapy might have been so successful. This patient was treated with several phage cocktails that were given sequentially, leading to rapid sequential development of bacterial resistance [20]. Most importantly, we find that the level of resistance increases rapidly when sensitive bacteria decline due to the (unsuccessful) phages in the first cocktail, and that this may markedly reduce the efficacy of subsequent cocktails. Thus, similar to lessons learned from treating rapidly evolving viruses like HIV, we find that it is important to ‘hit hard and early’ [13] by providing all phages at the same time, and to increase the ‘genetic barrier’ [2] by providing as many phages as possible.

## 2 Results

The killing of bacteria by neutrophils has been studied widely *in vitro* [18, 19, 21, 30] and *in vivo* [9, 10]. Importantly, these data reveal that there are two types of thresholds, one for the neutrophils and one for the bacteria. The control of bacterial growth requires a minimum number of neutrophils, which has been called the critical neutrophil concentration (CNC) [18, 19, 30]. The *in vitro* data in these papers have been modeled successfully with exponential growth of bacteria, *B*, and mass-action killing by neutrophils, *N*, i.e., d*B/*d*t* = *rB* − *kNB*, revealing that the neutrophil density at which d*B/*d*t* = 0, i.e., 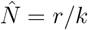, defines a CNC that is independent of the bacterial density [18, 19, 30]. Conversely, Drusano *et al*. [9, 10] inoculated the thighs or lungs of mice with increasing doses of bacteria, and found a threshold in the initial bacterial density above which the infection grows uncontrolled (one could call this a critical bacterial concentration, CBC). A CBC is a natural outcome of a model in which the number of bacteria that can be killed by the maximum number of neutrophils approaches a maximum [14, 23]. For instance, a model with a saturated killing rate, d*B/*d*t* = *rB* − *kNB/*(1 + *B/h*),in which a neutrophil can maximally kill *kh* bacteria per unit of time, has a CBC of 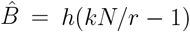. Since this saturation model approaches the mass-action model at low bacterial densities, i.e., when *B* ≪ *h*, this model also has a CNC at low bacterial densities. Malka *et al*.[21] vary both neutrophil and bacterial densities in *ex vivo* experiments, and find evidence for a double saturation model, d*B/*d*t* = *rB* − *kNB/*(1 + *B/h*_*k*_ + *N/h*_*N*_), which also obeys mass-action kinetics at low densities, but decreases the killing rate per neutrophil as a function of the neutrophil density (for which there is additional evidence [15]). However, for reasons of simplicity, we here adhere to the conventional model in which the killing rate only saturates with the bacterial density.

### 2.1 Results: mouse models

Leung and Weitz [15] developed a model allowing for a critical bacterial concentration, CBC, by also using the conventional saturated killing term, and in a second paper Roach *et al*.[26] showed that this model can explain how phage therapy can cure mice suffering from a rampant bacterial infection in the lungs by bringing the infection below the CBC. Because this model accounts for most of the *in vivo* observations during successful phage therapy in mice [26], we take their model as a starting point. The model defines phage-sensitive bacteria, *S*, phase-resistant bacteria, *R*, and bacteriophages, *P*,

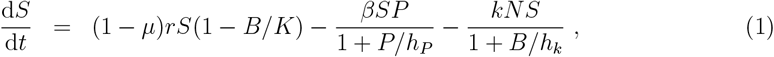

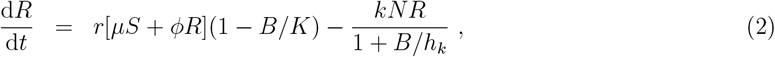

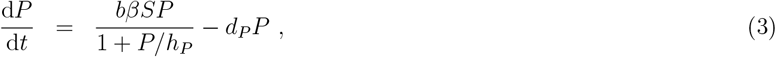

where *N* is the density of neutrophils, and *B* = *S* + *R* is the total density of bacteria. Bacteria grow logistically, with a net replication rate, *r*, have a carrying capacity, *K*, and a mutation rate, *µ*. Resistant bacteria may suffer from a fitness cost, *ϕ* ≤ 1. Mutations are not reversible, i.e., sensitive bacteria mutate into resistant bacteria, but mutations of resistant bacteria back to sensitive strains are not considered in the model (other mutations are included in the net replication rate). Time units are per hour and densities are defined in terms of numbers per gram of tissue [26]. At low phage densities the infection rate obeys mass action kinetics, where individual susceptible bacteria are infected at a rate *βP*. At high phage densities, i.e., when *P* ≫ *h*_*P*_, sensitive bacteria are infected at a maximal rate *βh*_*P*_, and when *P* = *h*_*P*_ the infection rate is half-maximal. The neutrophil killing rate was also defined by a conventional saturated killing term [15], saturating as a function of the bacterial density. At low bacterial densities the killing rate obeys mass action kinetics, with individual bacteria being killed at a rate *kN*, i.e., proportional to the density of neutrophils. At high bacterial densities individual bacteria are killed at a rate *kh*_*k*_*N/B* (which is proportional to the effector:target ratio, i.e., *N/B*). At very high bacterial densities, i.e., when *B* ≫ *kh*_*k*_*N*, the killing rate by neutrophils will therefore have a negligible effect, and bacterial densities will approach their carrying capacity, *K*. Phages are cleared at a *per capita* rate *d*_*P*_ (which is independent of their absorption to bacteria), and *b* defines a burst size, i.e., the number of phages produced due to the infection of a susceptible bacterium. The latter can be seen by inserting an equation for the infected cells, i.e.,

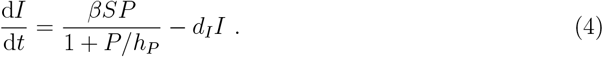

Then write

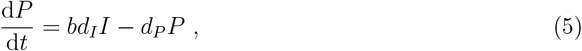

where *d*_*I*_ is the rate at which infected cells burst. The infected cell equation can subsequently be removed by the quasi steady state assumption (QSSA), d*I/*d*t* = 0, giving

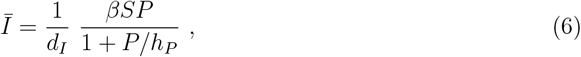

which when substituted into d*P/*d*t* in Eq. (5) yields Eq. (3) with a burst size *b*.

We continue by extending the model. Lytic phages typically take about half an hour to kill the bacteria they infected and produce a new generation of viral particles [22, 35]. Having about two generations per hour, and a burst size of approximately *b* = 100 particles [26], an upper bound for the replication rate of phages would be that all *b* = 100 phages produced by an infected cell are infectious, and together rapidly infect a total of a hundred bacteria, which after half an hour each produce another generation of *b* = 100 phages. If this were a synchronous process during the first few generations, we can index the number of infected cells for each half hour, i.e., *t* = 0, 0.5, 1, 1.5, …, by the map

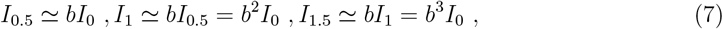

and so on. Thus, starting with one infected cell, *I*_0_ = 1, and ignoring viral clearance, there would be *I*_1_ = *b*^2^ = 10^4^ phages after one hour. To translate this level of expansion into the differential equations we are using, we can solve for the initial replication rate, *ρ*, from *I*(*t*) = *I*(0)e^*ρt*^, i.e., from 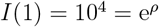 giving an upper bound *ρ* = 9.21 h^−1^.^1^

For a bacterial population at carrying capacity, *S* = *K*, the initial replication rate in the Roach *et al*. [26] model follows from Eq. (3),

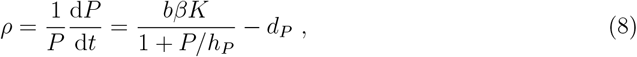

which at low phage densities would approach the unrealistically fast *ρ* = *bβK* − *d*_*P*_ = 5.4 × 10^4^ h^−1^ (parameter values are provided in the caption of Fig. 1). However, because the mice were treated by administering a large number of phages before the bacteria approached their carrying capacity (see Fig. 1), the initial *per capita* replication rate of the phages remained reasonably low, i.e., in Eq. (3) *P/h*_*P*_ ≫ 1 and *S* ≪ *K*. Since we consider treatments of rampant human bacterial infections in the second half of this paper, where the bacteria have approached their carrying capacity, we limit the initial replication rate by modifying the model in two ways. First, we slow the initial replication of the phages by not making the QSSA performed in Eq. (6), which limits the initial replication rate by the time scale at which bacteria are lysed (*d*_*I*_ = 2 h^−1^ [22, 35]). Second, because at high bacterial densities the infection rate should no longer be limited by bacteria, we modify the infection term into a double saturation function

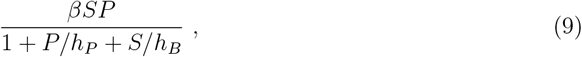

which naturally approaches *h*_*B*_*βP* when *S* → ∞, *h*_*P*_ *βS* when *P* → ∞, and a mass-action term *βSP* when both *S* and *P* are small [4, 7]. The model therefore changes into

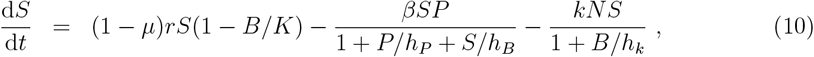

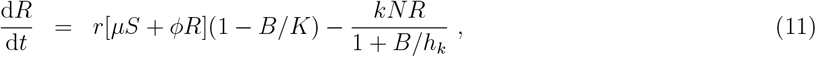

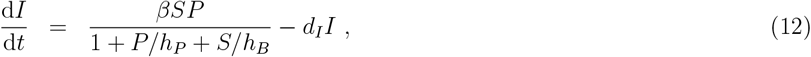

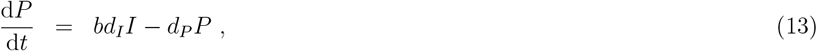

where Eq. (11) has remained the same as Eq. (2).

**Figure 1:**
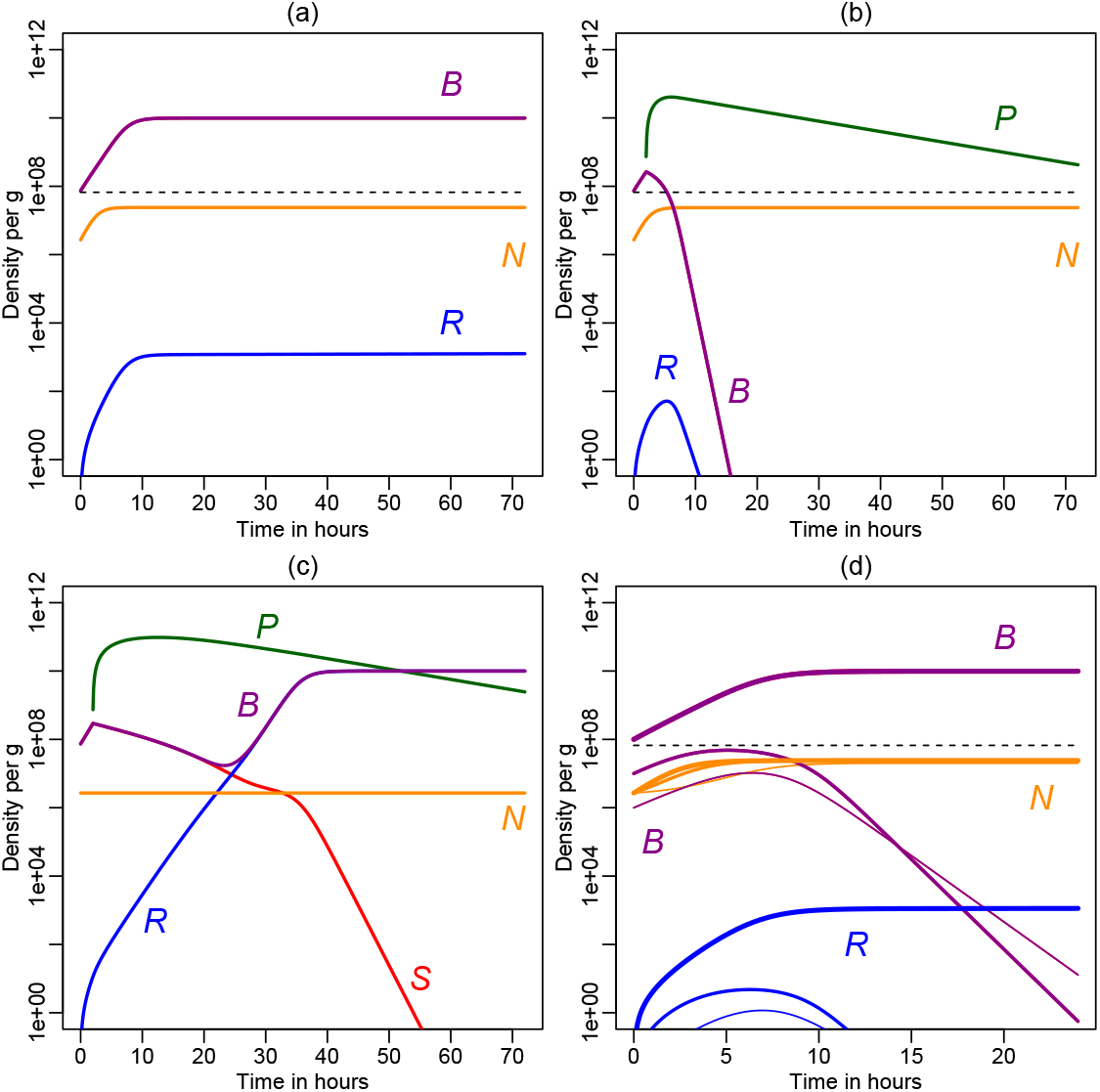
The characteristic behavior of the Roach *et al*. [26] model. Panels (a)–(c) illustrate their previous results with a large bacterial infection, *S*(0) = 7.4 × 10^7^ colony forming units (CFU/g) and *R*(0) = 0. Panel (a) depicts an infection without phages. Panel (b) depicts the same infection treated with a large dose of phages, *P* (2) = 7.4 × 10^8^ plaque forming units per gram (PFU/g), given after two hours. Panel (c) depicts the failure of the same treatment when neutrophils cannot expand beyond their initial level. Red lines depict sensitive bacteria, *S*, blue lines resistant bacteria, *R*, purple lines total bacteria, *B* = *S* + *R*, green lines phages, *P*, and orange lines neutrophils, *N*. The horizontal dashed black lines depict the CBC threshold 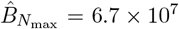 (this CBC is absent from Panel (c) because neutrophils cannot expand). Panel (d) illustrates that in the absence of phages bacterial infections starting below the CBC grow initially, and can only be controlled after the neutrophils have expanded significantly, and that bacterial infections starting above the CBC grow uncontrolled to their logistic limit. Infections are started with *S*(0) = 10^6^, 10^7^ to *S*(0) = 10^8^ CFU/g, and the line width of the curves identifies the three initial conditions. Roach *et al*. [26] parameters: *r* = 0.75 h^−1^, *K* = 10^10^ CFU/g, *ϕ* = 0.9, *µ* = 2.85 × 10^−8^, *β* = 5.4 × 10^−8^ g/(PFU h), *h*_*P*_ = 1.5 × 10^7^ PFU/g, *b* = 100, *d*_*P*_ = 0.07 h^−1^, *k* = 8.2 × 10^−8^g/(cell h), *h*_*k*_ = 4.1 × 10^7^ CFU/g, *r*_*N*_ = 0.97 h^−1^, *h*_*N*_ = 10^7^ CFU/g, *N* (0) = 2.7 × 10^6^ cells g^−1^ and *N*_max_ = 2.4 × 10^7^ cells g^−1^.

The initial replication rate of this model can be computed by considering the initial phase of a small population of phages (*P* ≪ *h*_*P*_) during a rampant bacterial infection (*S* = *K* ≫ *h*_*B*_), i.e., by simplifying Eq. (12) into d*I/*d*t* = *βh*_*B*_*P* − *d*_*I*_*I*. The initial phase of the infection is then described by the linear system,

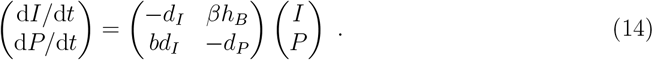

with eigenvalues

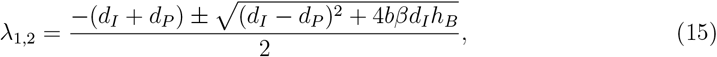

which has only one positive root defining the initial replication rate

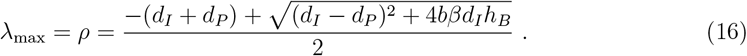

Solving for the new saturation constant, we find

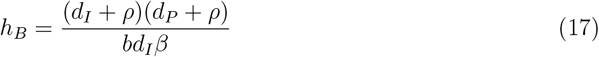

(See Gadhamsetty *et al*. [11] who used a similar approach for estimating the infection rate, *β*, during the acute phase of an HIV-1 infection). Substituting the parameters given in the captions of Figs. 1 and 2, and considering the upper bound *ρ* = 9.21 h^−1^, we find that in mice this replication rate is approached when *h*_*B*_ = 9.63 × 10^6^ CFU/g (which is similar to the other saturation constant, *h*_*P*_ = 1.5 × 10^7^ PFU/g, in Eq. (9)). Thus, if we were to use this saturation constant value for *h*_*B*_ in the double saturation function, Eq. (9), we are guaranteed to have a feasible initial phage growth rate in mice.

**Figure 2:**
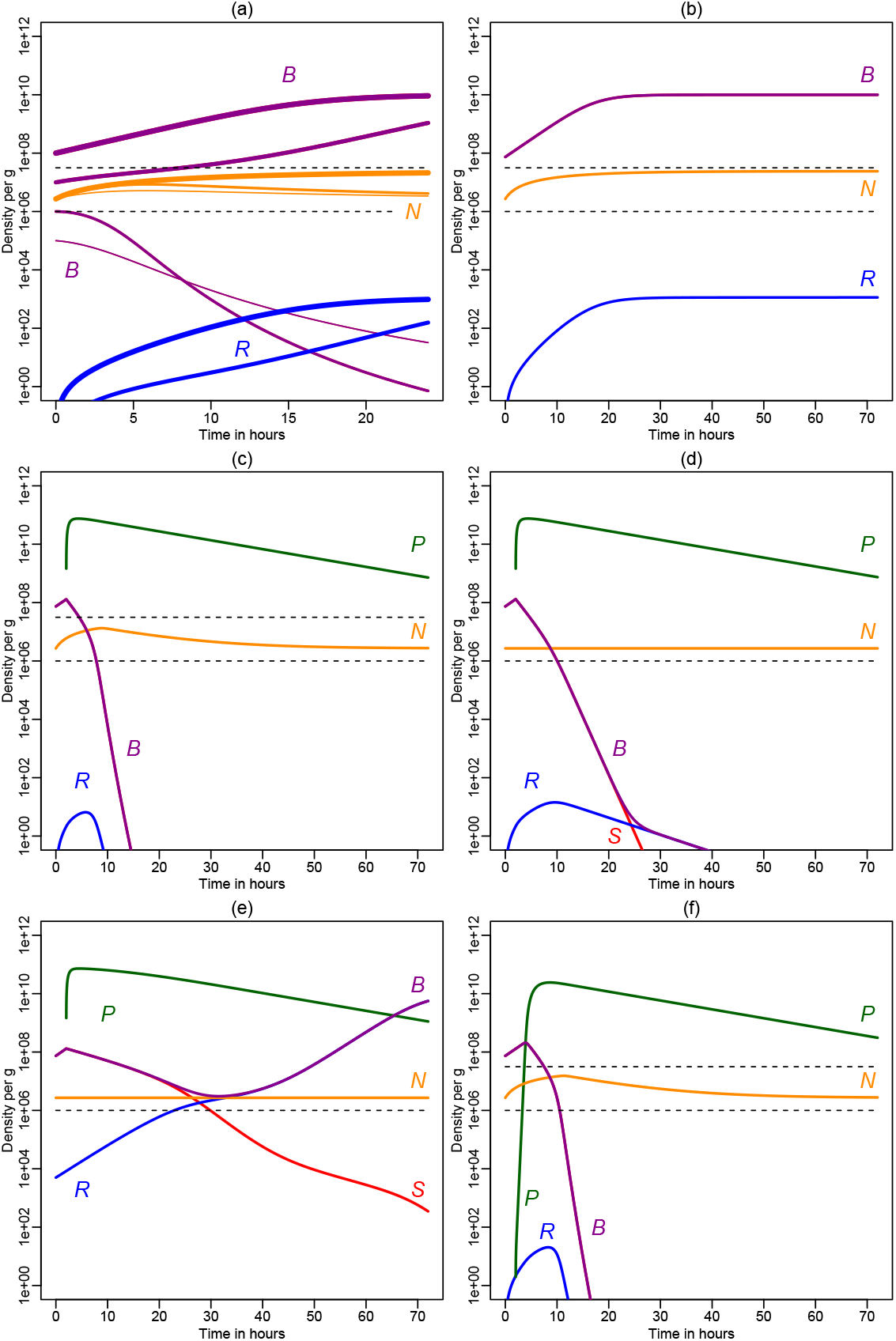
Similar behavior of the new model given by Eqs. (10 – 13) and (19). Panel (a) is like Fig. 1d with infections starting with *S*(0) = 10^5^, 10^6^, 10^7^ to *S*(0) = 10^8^ CFU/g. Infections starting below 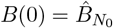 decline monotonically, and those above *S*(0) = 9 × 10^6^ CFU/g grow uncontrolled (in the absence of phages). Panel (b) is like Fig. 1a, an infection without phages. Panel (c) is like Fig. 1b, a controlled infection due to a large dose of phages, *P* (2) = 7.4 × 10^8^ PFU/g, given after two hours. Panel (d) depicts a controlled infection due to the same large dose of phages, in the absence of an increase in neutrophils (*α* = 0). Panel (e) depicts the evolution of resistance (like Fig. 1c) obtained by decreasing the infection rate 2-fold, and starting with *R*(0) = 5000 CFU/g resistant bacteria. Panel (f) is like Panel (c), a controlled infection, but due to a small dose of phages, *P* (2) = 1 PFU/g, given after two hours. The horizontal black dashed lines represent the low and the high CBC, 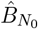 and 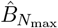, respectively (the high CBC is absent when *α* = 0). Colors as in Fig. 1, *S*: red, *R*: blue, *B*: purple, *P* : green, and *N* : orange. New parameters: *σ* = 2.25 × 10^5^ cells h^−1^, *d*_*I*_ = 2 h^−1^, *d*_*N*_ = 0.0833 h^−1^ (i.e., 0.5 d^−1^), *r* = 0.3 h^−1^, *k* = 1.5 × 10^−7^g/(cell h), *α* = 7.9, *d*_*N*_ = 1*/*12 h^−1^, *h*_*B*_ = 9.63 × 10^6^ CFU/g, *h*_*N*_ = 10^5^ CFU/g, and *h*_*k*_ = 2.86 × 10^6^ CFU/g.

#### Neutrophil recruitment

In the original model [15], the recruitment of neutrophils into the lungs was modeled by logistic growth at a maximum rate, *r*_*N*_, which saturated as a function of the bacterial density, *B*,

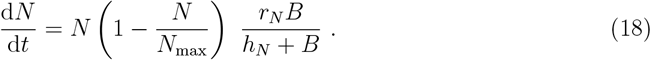

The carrying capacity, *N*_max_, of this logistic equation defined the maximum density of neutrophils per gram of tissue, and their initial density (in the absence of bacteria) was defined as an independent parameter, *N* (0) (note that in the absence of bacteria d*N/*d*t* = 0). The neutrophil equation was fitted to neutrophil dynamics data obtained from the lungs of mice stimulated with LPS [25], which suggested an initial density of *N* (0) = 2.7 × 10^6^ neutrophils/g, and maximal density of *N*_max_ = 2.4 × 10^7^ cells/g (i.e., an almost 9-fold increase). We modify the neutrophil model for several reasons. First, recruitment of neutrophils into tissues actually consists of two phases, i.e., a very fast phase where circulating neutrophils leave the blood locally to enter an inflamed tissue, and (2) a slower phase where the number of circulating neutrophils increases due to an increased release of immature and mature neutrophils from the bone marrow [1, 33]. The 24 h time scale of the Reutershan *et al*. [25] experiments, and the fact that the number of neutrophils in the lungs exceeded the initial number present in the circulation, implies that the data largely reflect an increased release from the bone marrow. Second, since in Eq. (18) d*N/*d*t* = 0 when *B* = 0, neutrophil densities fail to return to their initial density after a bacterial infection has been cleared. We therefore propose

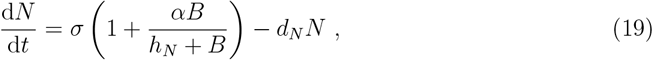

where *σ* is the normal production rate of circulating neutrophils by the bone marrow, and *α* ≃ 8 would allow this to be increased approximately 9-fold at high bacterial densities. The neutrophil death rate *d*_*N*_ allows the neutrophil density to return to the initial state *N* (0) = *σ/d*_*N*_, after clearing the bacterial infection. The maximum neutrophil density that is approached for large bacterial densities is *N*_max_ = *σ*(1 + *α*)*/d*_*N*_. Thus, we replace their logistic growth with Eq. (19) and adopt the previous parameters by defining, *σ/d*_*N*_ = *N* (0) and *α* = *N*_max_*/N* (0) − 1, such that we have the same initial and maximal density of neutrophils. The loss rate of the neutrophils, *d*_*N*_, defines the time scale at which the new *N*_max_ is approached, and we have tuned *d*_*N*_ and *h*_*N*_ to have a similar increase of neutrophils during the first phase of the infection. Below we will adapt this model for fitting human data. The more mechanistic Eq. (19) has the additional advantage that we have reasonable estimates for *d*_*N*_ in humans.

#### Neutrophil killing rates

The killing rates of the original model were based upon *in vivo* experiments in the lungs of mice in which bacterial expansion was studied, as a function of the number of bacteria inoculated [10]. This data revealed that initial bacterial densities smaller than *B*(0) = 10^7^ colony forming units per gram (CFU/g) can be controlled by the neutrophil response, and that larger densities grow uncontrolled [10]. By describing the data with a model with a saturated killing term similar to the one in Eq. (1), i.e.,

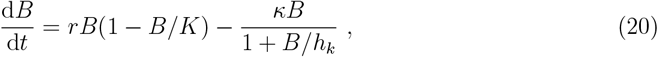

where we combine the killing rate and the neutrophil density into *κ* = *kN*, we have a model allowing for such a threshold [23]. This threshold can be found from Eq. (20) by computing the bacterial density at which d*B/*d*t >* 0, given a total neutrophil killing rate *κ*. For bacterial densities way below the carrying capacity, one finds that 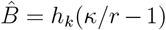 defines a CBC below which bacterial infections can be controlled. (Leung and Weitz [15] derive a similar threshold by solving the full quadratic equation defined by setting d*B* = 0*/*d*t* in Eq. (20)). Since *κ* = *kN*, this CBC depends on the neutrophil density, and because the model approaches two typical neutrophil densities, *N* (0) and *N*_max_, we also define two thresholds. The lower CBC threshold, 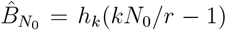, defines bacterial infections that are immediately controlled, and need no inflammatory response, and the upper CBC threshold, 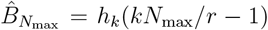, defines bacterial densities above which the maximum neutrophil response fails to control the infection. The two thresholds will be depicted as dashed or dotted lines in all figures.

Drusano *et al*.[10] fitted data they collected on the rate of *Pseudomonas aeruginosa* killing by neutrophils in a murine model of pneumonia with a mathematical model, and some of the parameters they estimated were adopted by Roach *et al*.[26], i.e., *r* = 0.75 h^−1^ (a doubling time of 0.9 h) and *κ* = 2 h^−1^. Assuming that this estimated total killing rate, *κ*, was due to a neutrophil density at its maximum, Leung and Weitz [15] subsequently estimated the mass action killing rate *k* = *κ/N*_max_ [26]; see Table 1. By finally setting the saturation constant to *h*_*k*_ = 4.1 × 10^7^ CFU/g, their model was given the property that bacterial infections starting at a density below *B*(0) ≃ 10^7^ CFU/g are controlled, whereas larger densities approach the carrying capacity (see Fig. 1d). For these parameters individual neutrophils can maximally kill *kh*_*k*_ = 3.4 CFU/g bacteria per hour, and the upper threshold is located at 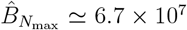 CFU/g (see Table 1). However, the combination of their bacterial replication rate, *r*, their neutrophil killing rate, *k*, and their normal neutrophil level, *N* (0) = 2.6 × 10^6^, does not allow small bacterial infections to be controlled without an inflammatory response, i.e., *kN* (0)*/r <* 1, implying that their lower threshold, 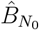, is negative, and that *N* (0) is below the CNC.

**Table 1:**
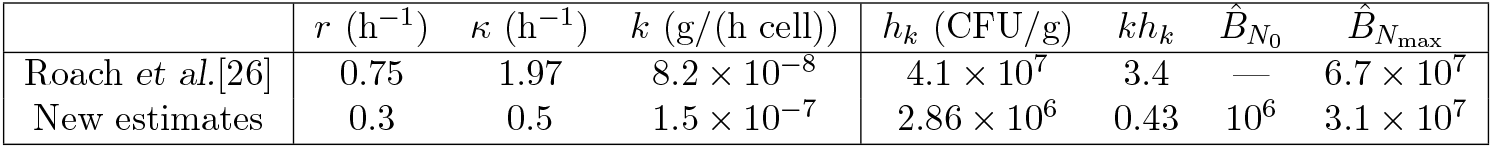
The killing parameters of Roach *et al*. [26], and new estimates obtained by refitting the Drusano *et al*. [10] experiments in mice using Eq. (20).

Unfortunately, these parameter estimates suffer from several uncertainties. First, Drusano *et al*.[10] actually fitted their data with a different model

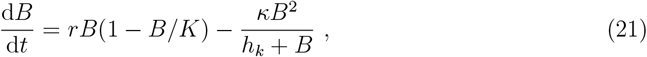

having a saturated *per capita* killing rate, rather than a saturated total killing rate. Second, the data they used was relatively sparse and only consisted of 5 data points, i.e., 5 pairs of an initial bacterial density, *B*(0), and a corresponding density, *B*(24), after 24 h (see the circles in Fig. S.1a). Apparently their fitting procedure suffered from a parameter identifiability problem, because the standard deviations of most parameter estimates are larger than their mean [10]. We have refitted the data with Eq. (20), while reducing the identifiability problem by (1) fixing the carrying capacity to the one estimated previously, and (2) by estimating the growth rate, *r*, from an experiment where Drusano *et al*. [10] depleted the neutrophils and observed bacterial densities to increase from *B*(0) = 3 × 10^4^ to *B*(24) = 4.7 × 10^7^ CFU/g (see the bullets in Fig. S.1a). Since the latter is well below the carrying capacity, *K* = 10^10^ CFU/g, we assume exponential growth, and solve for *r* from *B*(24) = *B*(0)e^*r*24^ giving *r* = 0.306 h^−1^ (i.e., a doubling time of 2.3 h). Fixing *r* and *K* we obtain the estimates *κ* = 0.5 h^−1^ and *h*_*k*_ = 1.5 × 10^7^ CFU/g (see Table 1 and the legend of Fig. S.1 for additional information). Note that, even after fixing *r* we find a strong negative correlation between *k* and *h*_*k*_ (not shown), suggesting that even the current estimates of *κ* and *h*_*k*_ are not reliable.

Finally, having somewhat better estimates for the total killing rate *κ*, we still need to estimate the killing rate per neutrophil, *k* = *κ/N*, as the neutrophil density should also be increasing in the Drusano *et al*. [10] experiments (i.e., *κ/N*_max_ *< k < κ/N* (0)). To estimate a new killing rate from our new estimate *κ* = 0.5 h^−1^ for *r* = 0.3 h^−1^, we first apply the constraint that small bacterial infections should be controlled at normal neutrophil levels, i.e., *kN* (0) *> r*, implying that *k > r/N* (0) = 1.1 × 10^−7^ g/(h cell). Setting *k* = 1.5 × 10^−7^ g/(h cell), which is almost 2-fold larger than the previous estimate, we next constrain the estimate of *h*_*k*_ from the fact that the lower CBC, 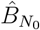, should be much smaller than 10^7^ CFU/g (as higher densities grow uncontrolled [10]). Solving *h*_*k*_ from 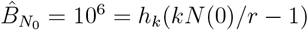 yields *h*_*k*_ = 2.86 × 10^6^ CFU/g, which translates into a maximum killing rate of *kh*_*k*_ = 0.43 CFU/g per neutrophil per hour. For these new parameter estimates the upper CBC becomes 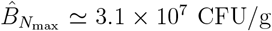 (which seems reasonable because this is just 2-fold lower than the previous estimate [26]; see Table 1). Summarizing, we obtain new parameter estimates from the same data, with a faster mass-action killing rate, *k*, and a lower saturation constant, *h*_*k*_, which yields the new feature that small bacterial infections can be controlled without increasing the neutrophil density, and maintains the previous feature that large bacterial infections cannot be controlled.

### 2.2 Results: mouse experiments

The most important contribution of the Roach *et al*. [26] paper was to demonstrate that the neutrophil immune response is essential to make phage therapy for *Pseudomonas aeruginosa* work in mice. Mice were infected intranasally with large doses of *P. aeruginosa*, and two hours later a large dose of a Pseudomonas phage was given intranasally. Time courses of bacteria and phages were followed over 72 hours. Importantly, the phage therapy was beneficial only when the mice had normal neutrophil numbers, and when these were able to respond to the infection [26]. We illustrate their results in Fig. 1a–c. A large bacterial infection that starts just above the CBC approaches carrying capacity in 10 h (Fig. 1a). The same infection is controlled by a high dose of phages administered after two hours, by bringing it below the CBC within ten hours (Fig. 1b). The same phage therapy fails when neutrophils cannot expand (Fig. 1c), resulting in a chronic infection with phage-resistant bacteria. These model predictions were elegantly confirmed by experiments, where the failure of the therapy in Fig. 1c was tested with MyD88-deficient hosts that lack immune cell activation and recruitment to the site of infection [26]. The fact that for the original parameters the model required expansion of the neutrophils to control even very small bacterial infections is illustrated in Fig. 1d, where we depict a series of bacterial challenges, from *S*(0) = 10^4^ to 10^8^ CFU/g, that all grow initially, and all evoke an increase in the neutrophil densities. Fig. 1d also confirms that challenges above *S*(0) = 10^7^ grow uncontrolled (despite the fact that this is below the CBC located at 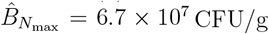; see the horizontal black line).

We have modified the Roach *et al*. [26] model by replacing the logistic growth term for the increase of the neutrophils by a more natural model having a source from the bone marrow (see Eq. (19)), by making the infection rate a saturation function of both the phages and the bacteria, by not making the QSSA for the infected bacteria, by increasing the killing rate, *k*, and by decreasing the saturation constant, *h*_*k*_, such that bacterial infections smaller than *B*(0) = 10^6^ CFU/g can be cleared without requiring increased production of neutrophils in the bone marrow. The latter implies that the new model has two CBCs, 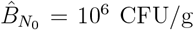 and 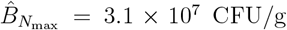, for the normal and the maximum number of neutrophils, respectively (see Table 1). The effect of the two thresholds is illustrated in Fig. 2a, where we depict a similar series of bacterial challenges, from *B*(0) = 10^5^ to 10^8^ CFU/g, in the absence of phages. Infections starting below 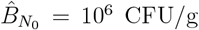 decline immediately, and infections starting with *B*(0) = 10^7^ CFU/g or more grow uncontrolled (similar to the original model).

Because we have tuned the parameters of the new neutrophil model to that of the previous model, we observe very similar establishment of a chronic infection with a large challenge of bacteria (Fig. 2b), and a very similar successful phage therapy with a large dose of phages after two hours (Fig. 2c). The main novel feature of the new model is that phage therapy can also work when the bacterial densities drop below the lower CBC; see Fig. 2d, where we have set *α* = 0 to prevent expansion of the neutrophils. Increased production of neutrophils is therefore not strictly required to make phage therapy work. The new model can also account for a similar failure of phage therapy, and the evolution of resistance (Fig. 2e), when we increase the initial number of resistant bacteria and decrease the infection rate. Finally, we observe in the new model that the initial dose at which the phages are administered hardly matters because they expand vigorously in the presence of a large density of susceptible bacteria (compare Fig. 2c with Fig. 2f), despite the fact that we have bounded the maximal replication rate by not making the QSSA d*I/*d*t* = 0, and by using the new infection term of Eq. (9). Likewise, replacing the single dose of phages with a small influx starting after two hours, representing the intravenous administration used to treat patients [8, 24, 28], yields a time course that is very similar to that shown in Fig. 2f (see below).

What have we learned from modeling phage therapy in mice? First, we confirm that the dose at which phages need to be administered to successfully control a bacterial infection hardly matters [15]. If the phages are able to infect the bacteria they will expand rapidly in a few generations, even in a model where we limit the replication rate of phages to *ρ* = 9.21 h^−1^. Clinically this is interesting because it is important that the phage preparations are sterile, and have endotoxin levels below FDA-approved levels [12, 24], which is easier to achieve with small than with large doses. Second, by re-estimating the rate at which neutrophils kill bacteria, we now have two thresholds in the model, and successful phage therapy no longer requires an increased neutrophil production since the bacterial infection will also become controlled when bacterial densities drop below the level that can be controlled by normal (non-inflamed) neutrophil densities. This could also be important clinically because it is not known how long neutrophil responses remain elevated during chronic bacterial infections. As a next step we develop this mouse model further by applying it to the human situation where patients are treated with cocktails of phages that the bacteria can independently evolve resistance to.

### 2.3 Results: a general human model

Because patients are treated with cocktails of phages, and bacteria independently evolve resistance to all of them, we have to extend the model by allowing for several strains of bacteria, and define the mutation rates at which these strains evolve from one another. Because patients are infused with phages we also add a time dependent source, *V* (*t*), to the ODE for the phages. We also add a strain-specific loss of bacteria to model treatment with antibiotics, because patients are often treated with various antibiotics, as sensitive and resistant strains frequently differ in their sensitivity to antibiotics [24]. For instance, Chan *et al*. [6] used phages binding the membrane proteins of a multi-drug efflux system, such that bacteria developing phage-resistance by down regulating these receptors became more sensitive to antibiotics. Because the concentration of phages in human serum can decline quite rapidly [28], we estimate the loss rate of phages, *d*_*P*_, by modeling the pharmacokinetics of the phages in serum after intravenous administration, as measured by Schooley *et al*. [28] (see Section S.2.3). Fitting simple models to that data we estimate that in humans *d*_*P*_ = 3.5 h^−1^ (see Fig. S.6). An alternative would be to allow for a loss of phages by absorption to bacteria, and/or a loss of phages due to the (slow) development of phage-specific antibody responses by the host [24]. Finally, since one of the potential mechanisms underlying the success of phage therapy is a loss of bacterial fitness due to their evolution of phage resistance, we extend the logistic bacterial growth model by including a density independent death rate, *d*_*B*_, of the bacteria, i.e., d*B/*d*t* = *ϕrB*(1 − *B/K*) − *d*_*B*_*B*. This allows the steady state of a bacterial strain, 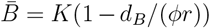, to depend on its fitness *ϕ*. In the previous model the steady state of bacteria in the absence of phages and neutrophils was defined as 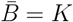, and it did not depend on their fitness *ϕ*.

To model bacterial strains resistant to several phages we will use a conventional bit string notation, where the strings ‘000’, ‘001’, ‘010’, ‘011’, ‘100’ have 0, 1, 2, 3, and 4 as their decimal interpretation, respectively. Also ordering the phages from right to left in the string, we define bacteria *B*_1_ with bit string *s*_1_ = 00 to be susceptible to phages *P*_1_ and *P*_2_, bacteria *B*_2_ with the bit string *s*_2_ = 01 to be resistant to phage one only, bacteria *B*_3_ with the bit string *s*_3_ = 10 to be resistant to phage two only, and bacteria *B*_4_ with bit string *s*_3_ = 11 to be resistant to both phages *P*_1_ and *P*_2_ (i.e., the bacterial strain number *i* is one plus the decimal interpretation of its string). A model with *n*_*P*_ different phages, each requiring a unique resistance mechanism, therefore needs bit strings of *n*_*P*_ bits defining 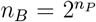 different types of bacteria, and *n*_*P*_ *n*_*B*_ types of infected cells, and a *n*_*P*_ × *n*_*B*_ susceptibility matrix *M*_*ij*_ defining all possible routes of infection. If a particular mutation were to confer resistance to subset of the phages, we combine these phages into ‘phage groups’ requiring a single bit in the bit string notation.

Since each phage group requires it own unique resistance mechanism, evolving resistance to them requires different ‘mutations’, i.e., deletions, insertions and point mutations, with different mutation rates. We therefore generalize the single mutation rate, *µ*, into a vector, *µ*_*i*_, where *i* = 1, 2, …, *n*_*P*_. Since each bacterial strain *i* can evolve from any other strain *j* we combine these mutations into a matrix, 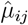. To keep the model simple we also only consider ‘gain of resistance’ mutations [15, 26], i.e., the upper diagonal of the mutation matrix is zero. For example, a fully susceptible *B*_00_ strain will mutate to *B*_01_ and *B*_10_ strains, at rates *µ*_1_ and *µ*_2_, respectively, and into strain *B*_11_ at a rate *µ*_1_*µ*_2_. For these four bacterial strains the mutation matrix would look like

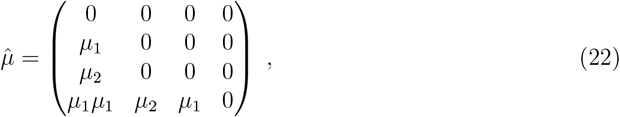

where each column defines the rate at which *B*_*i*_ mutates into the *B*_*j*_s, which is a loss term for *B*_*i*_ and a gain for a *B*_*j*_ (see below). Following the original model developed for mice [15, 26], mutations occur when bacteria divide. In chronically infected patients resistant bacteria are expected to be present before phages are introduced, as bacteria have evolved rapid resistance mechanisms over millions of years of co-evolution with phages [3]. The fitness, *ϕ*_*i*_, of bacterial strain *i* can be estimated from data (see Section S.2.2), or can be defined as a weighted sum (or a product) of the costs associated with each resistance mechanism, e.g., 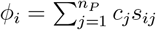, where *c*_*j*_ is the cost associated with mutation, *µ*_*j*_, and *s*_*i*_ is the bit string of bacterial strain *i*, and *j* is the phage number (and the position in the string when counting from right to left). Finally, bacteria may become more sensitive to killing by neutrophils when they become resistant by modifying their outer membrane, [20]. We allow for this by giving each bacterial strain its own rate, *k*_*i*_, of being killed by neutrophils.

Since we now consider several phages we need to reconsider the saturation in the infection term. As different phages, *P*_*i*_, may use the same receptor molecule on the bacteria to enter and infect, we mechanistically derive a high-dimensional infection term by ordering the phages into groups using the same receptor, and deriving a saturated infection term for all phages within one group (see the cartoon (Fig. S.4) and the full derivation in Section S.2.1). Due to its complexity we propose to lump all phages within one group into one “super phage” with an “average” infection rate, *β*_*i*_, and saturation constant, *h*_*P*_. This has the additional advantage that the model can be written with one phage-resistance mechanism per group, i.e., if the bacteria down regulate the receptor used by group *i*, all phages in that group would be affected. Thanks to this simplification, the form of the infection term remains similar to the one used previously [26].

Summarizing, we consider a population of sensitive and resistant bacteria, *B* = ∑_*i*_ *B*_*i*_, growing logistically. Bacteria, *B*_*i*_, are infected by different groups of phages, *P*_*j*_, to form infected cells, *I*_*ij*_, which burst and produce more phages, *P*_*j*_. Bacteria are killed by a population of neutrophils, *N*, and the rate of killing is half-maximal when *B* = *h*_*k*_. Bacteria can only be infected by the phage groups they are susceptible to, which is summarized in the susceptibility matrix *M*_*ij*_ defining all possible routes of infection. All of this can be formalized into the following model:

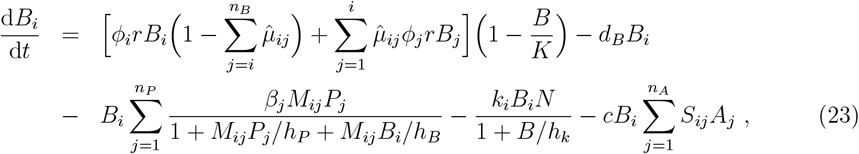

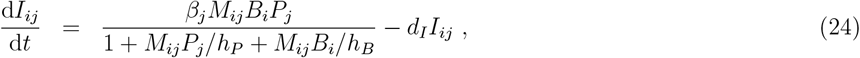

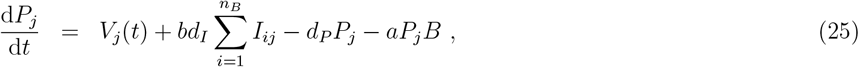

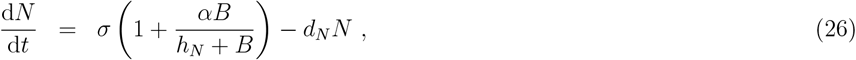

for *i* = 1, 2, …, *n*_*B*_ and *j* = 1, 2, …, *n*_*P*_, and where *n*_*P*_ defines the number of phage groups. The vector *β*_*j*_ defines the infectivity of phage strain *j, M*_*ij*_ defines which bacterial strain is resistant (*M*_*ij*_ = 0) or sensitive (*M*_*ij*_ = 1) to what phage, and *n*_*A*_ is the number of potentially effective antibiotics a patient is treated with, where *A*_*j*_ is the concentration of the *j*^th^ antibiotic, and *S*_*ij*_ defines the sensitivity of bacterial strain *i* for antibiotic *j*. Eq. (25) for the phages is extended with a time-dependent and phage-specific intravenous infusion rate, *V*_*j*_(*t*), and a mass-action absorption to all bacteria at a rate *a*. (The latter is a simplification because bacteria sometimes down-regulate the receptor molecule that phages use for binding and entering as a resistance mechanism.)

#### Human parameter values

The original model was parametrized on mouse data per gram of tissue, and since bacterial and phage densities per gram of tissue are probably similar in mice and men, we use most parameters of the mouse model for the human situation (except when human estimates are available). Since patients with rampant bacterial infections typically have several tissues infected with bacteria, the neutrophil equation of the model can best be interpreted as the number of neutrophils in the circulation and peripheral tissues. In the absence of inflammation we define a normal value, *N* (0), and we allow this to be increased 10-fold at high bacterial densities, by setting *α* = 9. Because we use the faster loss rate of phages that we estimated for humans, *d*_*P*_ = 3.5 h^−1^ (see Section S.2.3), the phage densities now decline to much lower levels when the bacteria become resistant or disappear due to successful therapy. To allow a population of phages to go extinct (and not grow back from unrealistic ‘atto-fox’ levels when sensitive bacteria recover), we multiply the infection rate with a steep Hill function, i.e., we use a ‘truncated’ infection rate, *β*_*j*_*/*(1 +[*θ/P*_*j*_]^5^). Thus, the infection rate is halved when *P*_*j*_ = *θ*, it rapidly approaches zero when *P*_*j*_ *< θ*, and it rapidly approaches *β*_*j*_ when *P*_*j*_ *> θ*. Since we are modeling one gram of tissue, and one PFU should approximately correspond to a single phage, we set *θ* = 10^−4^ PFU/g to allow phages to go extinct when there is less than one particle in a total body weight of about 10 kg of relevant tissue (setting *θ* either 10-fold higher or lower does not affect our main results).

We estimate the infection rate of the various phages by the successfulness of the treatment. Phage therapy will be successful when it reduces the bacterial density several orders of magnitude, from its carrying capacity to breach a CBC below which neutrophils can eliminate the infection (this CBC may depend on the bacterial strain when strains experience different killing rates, *k*_*i*_). The bacteria will only decrease when the force of infection, *F*, exceeds the maximal net replication rate, *r* − *d*_*B*_. At high phage densities, i.e., *P*_*i*_ ≫ *h*_*P*_, the maximum infection rate per phage group approaches *β*_*i*_*h*_*P*_ (summing up to a force of infection 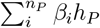 for all phages). Since the phages rapidly approach high densities, we define a critical infection rate above which one phage group can control the bacteria, by solving *β*_crit_*h*_*P*_ = *r* − *d*_*B*_, i.e., *β*_crit_ = (*r* − *d*_*B*_)*/h*_*P*_ = 2 × 10^−8^ g per PFU per hour. To maximize the initial replication rate of phages to *ρ* = 9.21 h^−1^, we now substitute the human parameters from Table S.2 into Eq. (17) and estimate that *h*_*B*_ = 3.56 × 10^7^ CPU/g (for *β* = *β*_crit_).

#### Single phage group

To explore the behavior of this new ‘human’ model we first simulate a simplified phage therapy with a single phage group, while varying the infection rate, *β*, and the mutation rate, *µ*. We consider a bacterial infection at steady state, with a corresponding increased number of neutrophils, and start with an arbitrary small number of phages, *P*_1_(0) = 1 PFU/g, on day zero (see Fig. 3). Because bacterial densities are high the phages approach their maximum replication rate of *ρ* = 9.21 h^−1^, and although their replication rate slows down when the phage densities approach *h*_*P*_, they very rapidly expand to densities exceeding 10^10^ PFU/g (Fig. 3). Because phages replicate so rapidly when bacteria are abundant, this confirms that the initial dose hardly matters.

**Figure 3:**
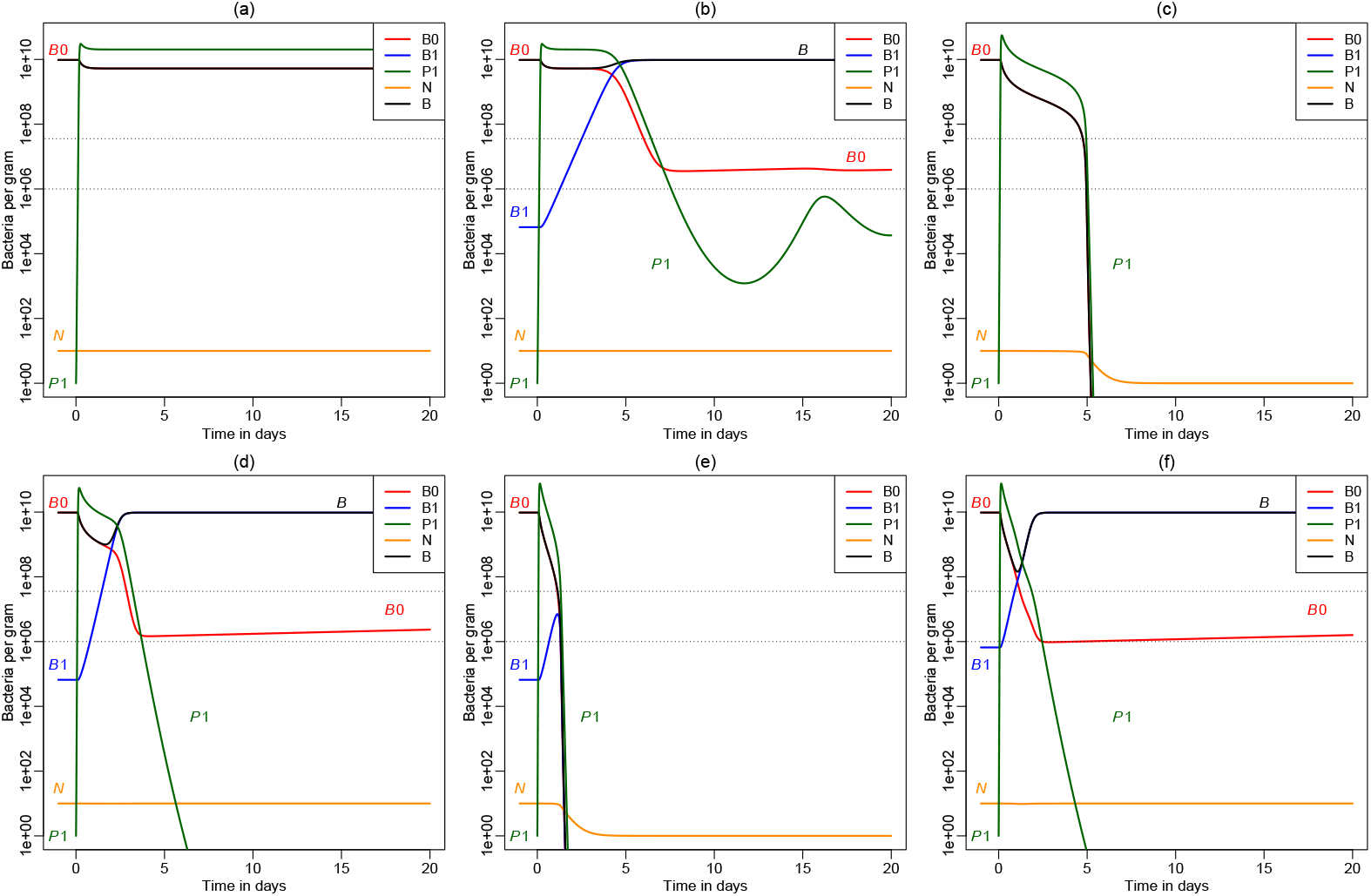
Exploring the human model with a single phage. Sensitive bacteria are now denoted as *B*_0_ and resistant bacteria as *B*_1_ (instead of *S* and *R* in the previous model). In Panels (a & b), (c & d), and (e & f), we set *β* = 10^−8^, *β* = 2 × 10^−8^, and *β* = 3 × 10^−8^ g per PFU per hour, respectively, i.e., below, at, and above *β*_crit_. In Panels (a) and (c) the mutation rate is set to zero, in Panels (b), (d) and (e) we have kept the *µ* = 2.85 × 10^−8^ h^−1^ that was used by Roach *et al*. [26], and in Panel (f) the mutation rate is increased 10-fold. In the absence of resistance, the phage clears the bacterial infection when *β* ≥ *β*_crit_; compare Panels (a) and (c). In the presence of resistance, an infection rate larger than *β*_crit_ need not be sufficient, because the resistant *B*_1_ bacteria may breach the upper CBC threshold 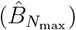; see Panel (d) and Panel (f). In Panel (b) the phages do not go extinct: the system slowly approaches a steady state dominated by a large population of resistant bacteria, with neutrophils at their maximum density, and the phages and the sensitive bacteria approach this steady state in a dampened oscillation (see a longer simulation in Fig. S.2a). In Panels (d) and (f) the phages are declining while the sensitive bacteria, *B*_0_, are slowly recovering. Here the phages do go extinct, and the sensitive bacteria slowly take over from the resistant bacteria due to their 10% higher fitness (see Fig. S.2b and Fig. S.2c). For better visibility the neutrophil concentration is divided by *N* (0) (i.e., *N* is scaled between one and ten). Novel parameter values: *ϕ*_1_ = 1, *ϕ*_2_ = 0.9, *r* = 0.31 h^−1^, *d*_*B*_ = 0.01 h^−1^, *d*_*P*_ = 3.5 h^−1^, *h*_*B*_ = 3.56 × 10^7^ CFU/g, *k*_1_ = *k*_2_ = 0.405 ^−1^h, *a* = 0, and 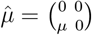; see Table S.2.

Phage therapies with phages having a low infection rate (i.e., *β < β*_crit_) fail because the sensitive bacteria just approach a somewhat lower level (see Fig. 3a; where there are no resistant bacteria because we set *µ* = 0), or only have a similar small transient effect because the resistant bacteria rapidly take over, and the bacterial infection returns to a similar high steady state (Fig. 3b). Since the phages are declining after most of the bacteria have become resistant, the sensitive bacteria start to recover slowly (from day 7 onward in Fig. 3b). Ultimately, this also allows the phages to recover, and the system slowly approaches a steady state dominated by resistant bacteria and high neutrophil levels, and about 10^7^ CFU/g sensitive bacteria maintaining a population of about 10^5^ PFU/g phages (Fig. S.2a). In the absence of resistance, phages with a higher infection rate of *β*_crit_ suffice to control the bacterial infection (Fig. 3c), but in the presence of resistance this infection rate need not be sufficient (Fig. 3d), because the bacteria rapidly become resistant and the phages go extinct (Fig. S.2b). Disturbingly, the ‘resistant’ steady state is approached faster when the phages are more infectious, because phages having a higher infection rate lead to a steeper decline of the bacteria, which reduces the logistic competition, and allows the resistant bacteria to replicate faster (compare the total bacterial load, *B*, in Fig. 3b with that in Fig. 3d).

Increasing the infection rate further allows the phages to clear the bacterial infection in the presence of resistance (Fig. 3e), because more infectious phages more rapidly deplete the sensitive bacteria, which here brings the total bacterial density below the upper threshold CBC, 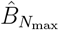, before the resistant bacteria have taken over. Note that resistant bacteria start declining when the total, *B*, breaches the upper threshold, that the phages disappear afterwards decline quite rapidly due to their relatively high turnover, and that the neutrophils return to their normal steady state (Fig. 3c and e). However, a therapy with the same more infectious phages fails when the bacteria more easily become resistant (Fig. 3f), because it then takes less time for the resistant bacteria to breach the upper threshold CBC.

Summarizing, whether or not phages can control a rampant bacterial infection depends both on their infection rate and on the rate by which the bacteria develop resistance, as phage therapy will fail as soon as the resistant bacteria breach the upper CBC. Although it should always be better to treat with highly infectious phages, an important new lesson learned is that resistant bacteria expand more quickly when the phages are more infectious. Having a basic understanding of the new model we next use it to simulate phage therapy with two phages.

#### Two phage groups

Consider a phage therapy consisting of two phages (or two cocktails of different phage groups) that are given sequentially. For simplicity, we consider phage groups with identical infection rates, and bacterial strains with identical mutation rates, killing rates, and fitness costs (see Fig. 4). Since a therapy with a single phage failed because of rapid development of resistance (Fig. 3c), we add on a second phage group on day five (Fig. 4a), or day two of the treatment (Fig. 4b and c). The former fails because the frequency of bacteria resistant to both phage groups (*B*_11_ denoted by orange lines), markedly increased when the bacterial strains resistant to the first group (*B*_01_ denoted by blue lines) took over. This is a natural result because fully resistant bacteria are just one mutational step away from bacteria resistant to the first cocktail, and two steps away from susceptible bacteria (see Eq. (22)). One can prevent double-resistant bacteria to breach the upper CBC by giving the second treatment earlier (e.g., at day two; Fig. 4b), which makes the combined treatment successful. Obviously, this timing depends on the initial frequencies of resistant bacteria (see Fig. 4c where we increased the mutation rates 10-fold), and the optimal treatment would be to give both phage groups at the same time (not shown).

**Figure 4:**
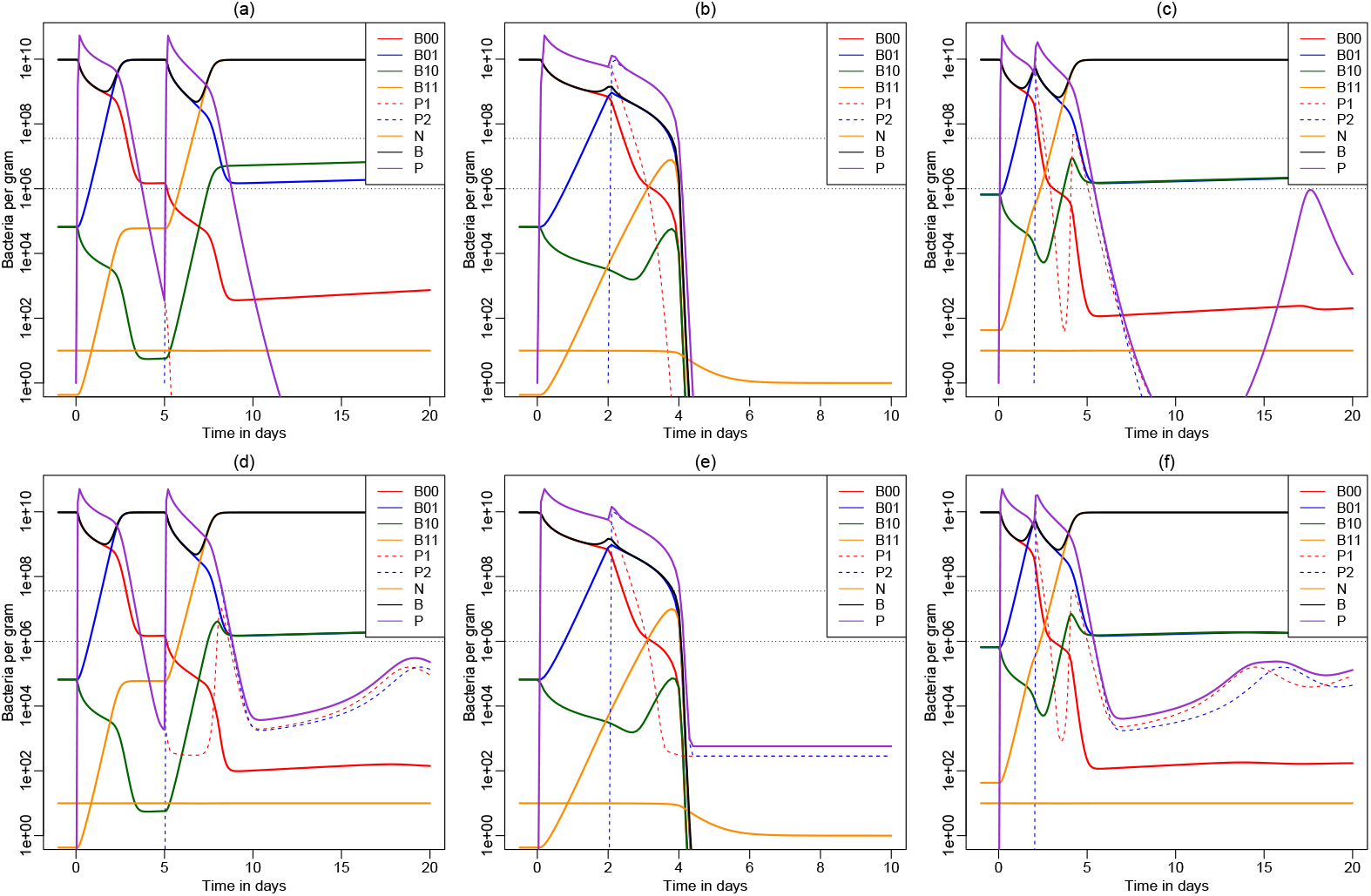
Simulated phage therapy with two phages administered sequentially. The second phage is either given fairly late (at day five (Panels a and d)) or early (at day two: Panels b, c, e & f). Sensitive bacteria are now denoted as *B*_00_, and the 3 types of resistant bacteria as *B*_01_, *B*_10_ or *B*_11_, respectively. Both phages have the critical infection rate *β* = 2 × 10^−8^ g/(PFU h). In the upper row the phages are given as a small single shot at day 0, 2 or 5, i.e., *P*_1_(0) = 1 and *P*_2_(2) = 1 or *P*_2_(5) = 1 PFU/g, while in the lower row the treatment is given in the form of a continuous infusion, i.e., 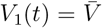 at all time points, and *V*_2_(*t*) = 0 at the start until we set 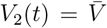 when *t >* 5 days (Panel d) or *t >* 2 days (Panels e & f). The first five days in Panel (a), and the first two days in Panel (b) are therefore identical to the unsuccessful therapy shown in Fig. 3d. Adding a second therapy at day five fails to prevent the evolution of resistance (Panels a & d), starting earlier makes the same therapy successful (Panels b & d), but a 10-fold higher mutation rate again leads to failure due to the evolution of resistance (Panels c & f). Importantly, there is hardly any difference in the total bacterial load, *B*, between single shots of the phages (Panels a – c), and a therapy comprised of continuous infusion (Panels d – f). Phages do not go extinct in Panel (c), and not when they are administered by continuous infusion (see Fig. S.3c–f). Novel parameter values: 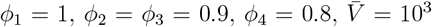 and *µ*_1_ = *µ*_2_ = 2.85 × 10^−8^ h^−1^ in the matrix shown in Eq. (22).

This resembles the ‘hit hard and early’ concept from antiretroviral therapy for HIV [13], where combinations of drugs requiring different escape mutations increases the genetic barrier to full resistance. A major difference is that phages are replicating, and that phage therapy therefore relies much less on the dosing schedule. This is illustrated in the bottom panels of Fig. 4 where we simulate a treatment consisting of a continuous infusion of the phages (rather than a small single ‘shot’ at a particular day). The high similarity in the total bacterial load, *B*, between the panels in the upper and lower row of Fig. 4 confirms that dosing makes little difference. Continuous infusion only prevents the phages from going extinct when the availability of sensitive bacteria declines (see Fig. S.3).

The most important new lesson learned is that phage therapies should be given as early as possible to slow down the evolution of full resistance. Continuous infusion may reduce the reversal of resistance in a bacterial population, but even this would not require high doses as phages start replicating as soon as susceptible bacteria reappear.

### 2.4 Results: the Patterson case

The first spectacular case of a patient recovering from a rampant bacterial infection with multidrug resistant (MDR) *Acinetobacter baumannii* after the administration of various cocktails of phages was described as the ‘Patterson case’ by Schooley *et al*. [28] and by Strathdee and Patterson [32]. Phage therapy was initiated about a 100 days after the initial infection while the patient was in coma, and although the bacteria developed resistance to all phages in the first two cocktails on a time scale of days, the patient woke up and started recovering [20, 28, 32]. Our aim here is to model this recovery during the first ten days of treatment.

The first two days of treatment used a cocktail of phages called ΦPC, that was given via the catheters draining the intra-abdominal abscess cavities. It remained unclear whether or not the phages were able to replicate in this fairly acidic environment, and the patient did not improve. A second cocktail consisting of four other phages (ΦIV) became available on day three, and because the two days of ΦPC treatment seemed to have little effect, the treatment was switched to dosing intravenously. After these four days of phage therapy the patient awoke. The therapy was stopped the next day because of clinical problems, and as soon as this was diagnosed to be due to an unrelated bacterial infection, the treatment was reinstalled two days later, and continued for more than a 100 days [28]. After a week, minocycline was added to the treatment, but because the bacteria later turned out to be resistant to minocycline [20] we do not include any antibiotics in our model (i.e., we set *c* = 0 in Eq. (23)). A third cocktail was started on day 113, and because we only model the first 10 days during which the patient recovered, this is also not considered. Both the intra-abdominal and the intravenous infusion is accounted for with the *V* parameter in Eq. (25), and this particular time schedule of the treatments is implemented by setting *V*_*i*_(*t*) = 0 on day 1 and 2 for the phages in the ΦPC cocktail, and on days 5 and 6 for both cocktails.

The genomes of the phages were sequenced and this revealed that they were all closely related to T4-like myophages, and that three out of the four phages in the ΦPC cocktail were actually identical [20]. Thus, the first cocktail consisted of two related phages called Maestro and AC4, while the second ΦIV cocktail consisted of four related phages called Navy1, Navy4, Navy71 and Navy97 (see Table 2). T4-like myophages use long tail fibers to bind to the phage’s receptor on the cell surface, and tail fiber analysis showed the myophages used in the cocktails had two different types of tail fibers, with Maestro, AC4, and Navy71 belonging to one group, and Navy1, Navy4, and Navy97 belonging to the other cluster. This agreed well with the observation that at day two, the bacteria became resistant to all three phages in the first cluster, and remained sensitive to the three in the second cluster (Table 2 in Liu *et al*. [20]). Importantly, this suggests that although the phages in the first ΦPC cocktail had little clinical effect, they were able to replicate and select for resistance. On day six, i.e., after four days of treatment, bacteria were resistant to all phages [20].

**Table 2:** Composition of the two cocktails used by Schooley *et al*. [28]. The 6 phages can be classified in two resistance groups by their tail fibers [20], which is indicated in the rows.

| Cocktail: | ΦPC | ΦIV |
| --- | --- | --- |
| Resistance cluster 1 | Maestro & AC4 | Navy71 |
| Resistance cluster 2 | — | Navy1, Navy4 & Navy 97 |

The fact that resistance comes in two clusters suggests that this can be modeled with two phage groups. Since Navy71, belonging to the first resistance cluster, was only present in the second cocktail (see Table 2), which was administered after the bacteria were already resistant to it, we did not model it separately. Because resistance to Navy1, Navy4 and Navy97 evolved simultaneously [20], we also treat the second cocktail as one uniform group of phages. Since bacteria became resistant to the phages in the second cluster by a common 6 base pair deletion in genes predicted to be involved in the synthesis of their capsule [20], a ‘mutation’ which led to the loss of the bacterial capsule should also make them resistant to the phages in first cluster. Thus, we ignore the *B*_10_ bacteria and consider only two ‘mutation’ rates in Eq. (22). The sequencing data confirmed that the resistant strains evolved from a common ancestor [20], and in the model we let sensitive bacteria develop resistance to cluster one at a rate 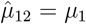, and let *B*_00_ and *B*_01_ strains shed their capsule at a rate 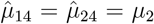. All other elements in the mutation matrix of Eq. (22) are zero (i.e., the *B*_10_ strain remains absent), and we will vary *µ*_1_ and *µ*_2_ to match the data.

In the first paper on the Patterson case [28], the development of phage-resistance was measured by culturing bacterial isolates from the patient in the presence and absence of the phages in the cocktails. We fit these *in vitro* data in Section S.2.2 to estimate the relative growth rates of the bacterial strains resistant to the first and second cocktail. The estimated fitness of pretreatment bacteria resistant to the first cocktail was estimated to be *ϕ*_2_ = 0.74, and their initial abundance was estimated to be high (21%) (see Table S.1). Culturing bacterial isolates taken at day six revealed that the estimated fitness of bacteria resistant to both cocktails is about *ϕ*_4_ = 0.33 (see Table S.1), with an estimated abundance of about 30% (at day six). The mutation rate leading to the *B*_01_ strain resistant to the first cocktail was set to *µ*_1_ = 0.05 to match the high initial abundance in the respiration experiments. The mutation rate leading to resistance to both cocktails was set to a 20-fold lower value to account for the later appearance of the *B*_11_ strains (that are also resistant to the first cocktail). Note that these mutation rates are orders of magnitude higher than those used above.

At the estimated abundance of about 20% resistant *B*_01_ strains in the pretreatment bacterial population, the resistant strains are way above the upper CBC (see day zero in Fig. 5a), and hence they rapidly take over during the first two days of treatment (whatever the infection rate of the phages). Because the first cocktail was administered into the fairly acidic environment of the abdominal abscess cavities, we assigned it an infection rate below the critical infection rate, i.e., *β*_1_ = *β*_crit_*/*2, and nevertheless observe that this unsuccessful cocktail does select for resistance in the bacteria. After two days of treatment the bacterial population changed from 81% susceptible *B*_00_ strains, to 98% *B*_01_ and 1% *B*_11_ strains (see day two in Fig. 5a). Because this 1% abundance of fully resistant *B*_11_ strains also exceed the upper CBC, treatment with the second cocktail would also be unsuccessful (whatever the infection rate of the phages); see Fig. 5a. Upon the introduction of the second ΦIV cocktail, the dominant *B*_01_ strains immediately decline, which accelerates the replication rate of the fully resistant *B*_11_ strains, that rapidly become dominant. Thus, the first cocktail markedly increases the resistance to the second cocktail even before this ΦIV cocktail is administered. Note that increasing the infectivity of the first cocktail would only speed up the development of full resistance, without making it successful.

**Figure 5:**
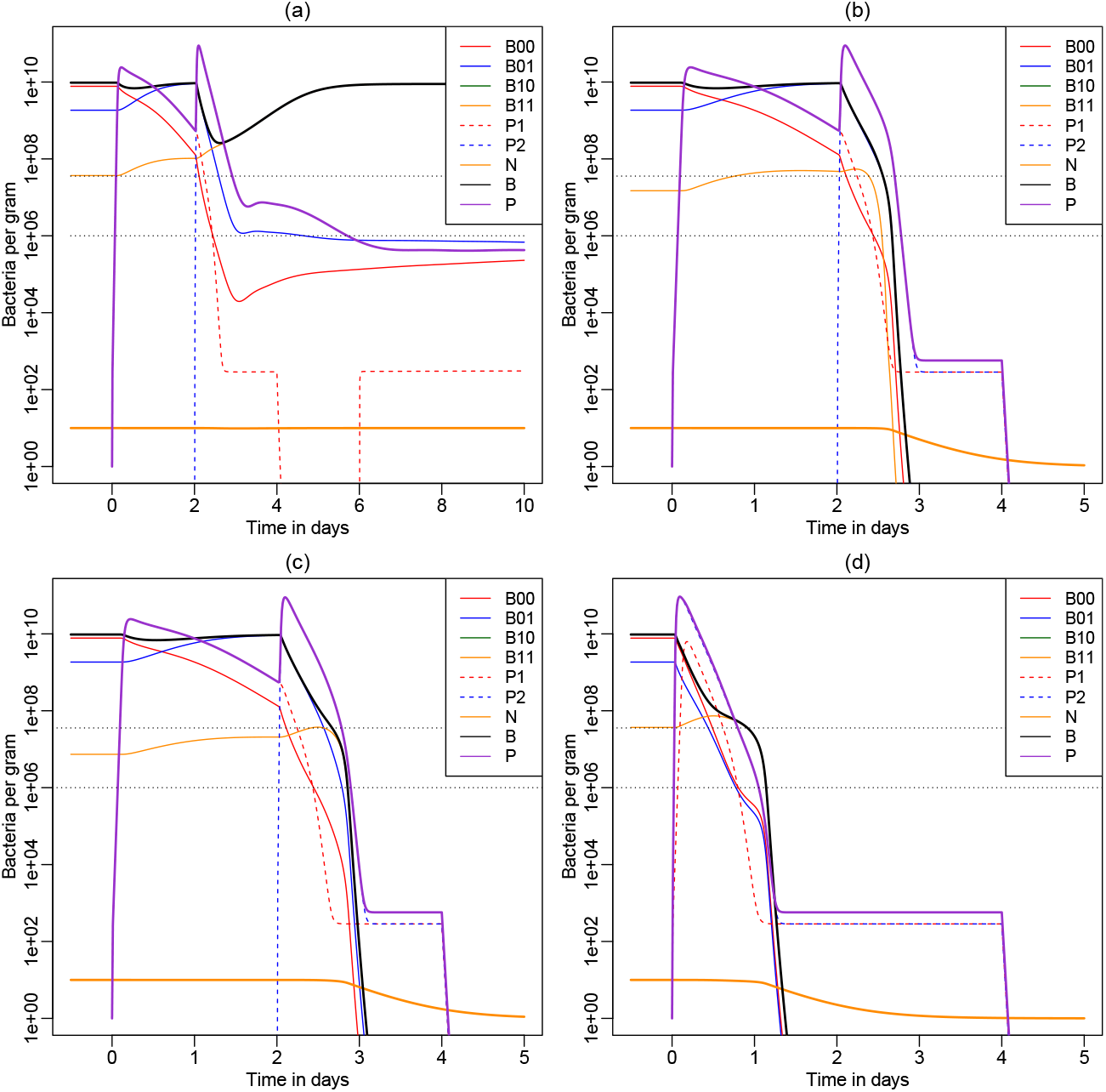
The Schooley *et al*. [28] patient modeled with two phage groups, i.e., the first cocktail given on day 1 (i.e., at *t* = 0) and a second given on day 3 (i.e., at *t* = 2). In Panel (a) the infection rate of cocktails is set to *β*_1_ = *β*_crit_*/*2 = 10^−8^ and *β*_2_ = 2*β*_crit_ = 4 × 10^−8^ g/(PFU h), which leads to failure of the treatment because at the estimated pretreatment level of resistance both types of resistant bacteria are above the upper CBC (with *µ*_1_ = 0.05 and *µ*_2_ = *µ*_1_*/*20 = 0.0025). In Panel (b) we allow for a 10-fold faster killing rate of the fully resistant *B*_11_ bacteria, which clears the infection. In Panel (c) we decrease the second mutation rate 5-fold (to *µ*_2_ = *µ*_1_*/*100 = 0.0005), which keeps the fully resistant bacteria below the upper CBC, which also makes the second cocktail successful. In Panel (d) the combination therapy of Panel (a) becomes successful when both cocktails are given at the same time, because the fully resistant bacteria have less time to expand. We use the same parameters as in Fig. 3 (see Table S.2), but use the estimated abundances and growth rates to set the mutation rates *µ*_1_ and *µ*_2_, and the fitness values *ϕ*_1_ = 1, *ϕ*_2_ = 0.74 and *ϕ*_4_ = 0.33 (see Table S.1). For better visibility the neutrophil concentration is divided by *N* (0) (i.e., *N* is scaled between one and ten).

Since Fig. 5a is not in agreement with the data as the second ΦIV cocktail was successful, we consider two possibilities. First, the fully resistant *B*_11_ strains are much more sensitive to being killed by neutrophils because they have a deficient capsule [20]. In Fig. 5b we therefore increase the killing rate of *B*_11_ bacteria, *k*_4_, by a factor 10, which is sufficient to explain the success of the treatment. Increasing the killing rate increases the upper CBC for fully resistant bacteria, which allows them to be controlled by neutrophils as soon as the total density of partially resistant bacteria has declined sufficiently by infection with the ΦIV phages (see Fig. 5b). Note that the increased killing rate also decreases the initial abundance of fully resistant bacteria (see day zero in Fig. 5b), which would be helpful if their abundance were to remain below the upper CBC.

A second possibility is that the initial abundances estimated from the respiration data are overestimated because (1) the relationship between the true bacterial density and the observed respiration need not be linear, (2) the background respiration that we subtracted from the data was large compared to the initial respiration of the bacteria, and (3) we do not have an estimate for the pretreatment abundance of the fully resistant bacteria because they hardly replicated in the respiration experiments [28]. Explaining the success of the second cocktail just requires a much lower abundance of the fully resistant bacteria, i.e., by setting *µ*_2_ = *µ*_1_*/*100, the fully resistant bacteria remain below the upper CBC, even after their rapid expansion during the first two days of treatment. This also suffices to explain the clinical success of the second cocktail (Fig. 5c).

Because the resistance to the second cocktail markedly increases during the treatment with the first cocktail, we improve the unsuccessful treatment of Fig. 5a by giving both cocktails at the same time (see Fig. 5d). Simultaneous administration makes the phage therapy successful because the fully resistant strain is not given the opportunity to expand during the treatment with the first cocktail. Summarizing, whether or not phage therapy can clear a rampant bacterial infection requires sufficiently low pretreatment levels of resistance, and administration of all phages at the same time may prevent the rapid expansion of fully resistant bacteria. Disturbingly, higher infection rates lead to a more rapid development of resistance.

### 2.5 Results: parameter sweep

The major advantage of an immediate parallel treatment with several phage groups is that the initial density of the fully resistant bacteria will still be at the pretreatment level, and that for each additional phage group requiring another resistance mutation, the expected pretreatment concentration will drop by its mutation rate. The lower the pretreatment concentration the longer it will take a fully resistant strain to breach the upper CBC, and hence the more likely that the treatment works by bringing the total bacterial load below the upper CBC. To address the question whether it is better to improve the infection rate of the phages, or to increase the diversity of a cocktail of phage groups (that are all given immediately), we performed a parameter sweep increasing 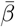 and *n*_*P*_ (see Fig. 6). Assuming identical mutation rates 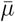 for all phage groups, we use Eq. (22) to define the full mutation matrix 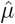. We vary the mutation rate in the range suggested by the cultured isolates, i.e., 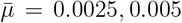 and 0.01 in Fig. 6a, b, and c, respectively, and we vary the common infection rate 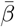128-fold by increasing it in 2-fold steps. This reveals that increasing the infection rate above *β*_crit_ hardly improves the successfulness of the therapy.^2^ Even extremely infectious phages require a minimum diversity of phage groups to make phage therapy successful at these pretreatment resistance levels.

**Figure 6:**
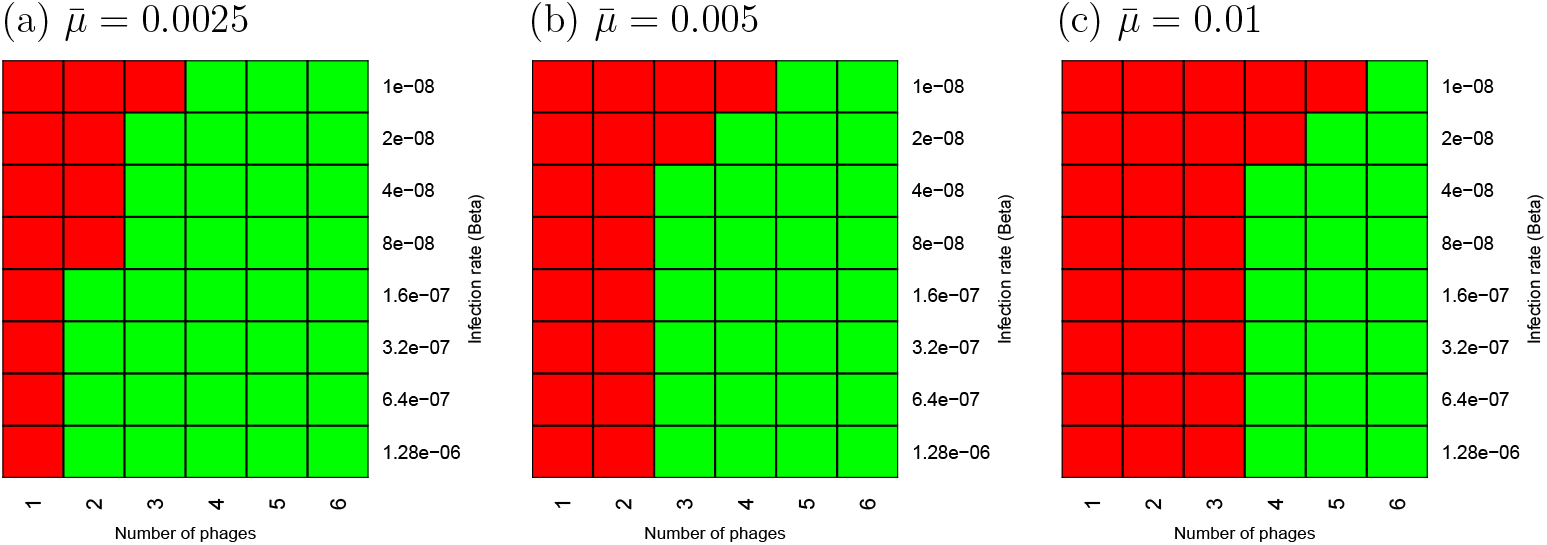
Parameter sweep, varying 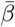 and the number of phage groups. In Panel (a), (b) and (c) we set 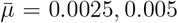 and 0.01, such that about 5%, 10% and 20% of the pretreatment bacteria are resistant to one of the phage groups, respectively. Each heat map depicts a parameter sweep, varying 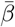 128-fold, and the number of phage groups from 1 to 6 (all start at *t* = 0). The fitness of the bacteria declines proportional to the number of resistance mutations of the bacterial strain, i.e., 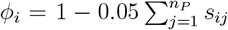, where *s*_*i*_ is the bit string of strain *i* and *j* denotes the position in the string. Green boxes denote successful therapy and red boxes depict failure. Whenever 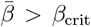, increasing the infection rate hardly affects the successfulness of the therapy. Even at very high infection rates, 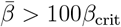 (where *β*_crit_ = 2 × 10^−8^ g/(PFU h), the pretreatment resistance levels, as set by the mutation rates, require a minimum number of phage groups, i.e., 1, 2 and 3, for 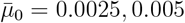 and 0.01, respectively.

## 3 Discussion

Having developed a novel mathematical model for phage therapy, and after confirming that this model can successfully describe clinical data, we now have an *in silico* tool enabling us to compare potential scenarios for treatment. Most importantly we have observed in our models that during the first days of successful treatment the replication rate of (partially) resistant bacterial strains markedly increases because they are released from the severe competition with the previously dominant pool of sensitive bacteria. Whether or not a therapy will be successful depends on time it takes resistant strains to breach the critical bacterial concentration (CBC) above which the immune system can no longer control the resistant bacteria. Unfortunately, the more infectious the phages in a first cocktail, the faster the release from competition, and the faster phage-resistant bacteria will breach this CBC. Indeed, once the infection rate of the phages suffices for controlling sensitive bacteria, a further increase of the infection rate is less important than extending the therapy with additional phages (Fig. 6). It is therefore better to provide all phages at the same time [20], to avoid this early expansion of phage-resistant bacterial strains. These findings mimic the lessons learned from antiretroviral therapy to HIV, where one combines several drugs, each requiring different resistance mutations [2], to hit the virus hard and early [13]. The main difference with antiretroviral and antibiotic therapy is that the dosing is less important in phage therapy because phages are rapidly ‘replicating drugs’ (the maximum replication rate of phages, *ρ* = 9.21 h^−1^, by far exceeds that of their bacterial hosts). This is clinically important because using lower doses would reduce the problem of residual endotoxin in phage preparations. Current recommendations of the Antibacterial Resistance Leadership Group (ARLG) in the United States are to use the highest safe and tolerated dose of bacteriophage products, although it is not clear that clinical outcome always improve with higher doses [24]. For instance, phage therapies in the United States typically use much higher doses that those in Europe, with similar clinical success rates between 55 and 70% [24]. Trials investigating the effect of dosage are ongoing [34], but the lowest doses are typically kept above 10^7^ PFU/ml. The main concern with using lower doses is that the phages may not reach the site of infection (which in our well-mixed model is never an issue). Nevertheless, because lower doses should reduce current problems with residual endotoxin in bacteriophage products, it would be interesting to further investigate this experimentally and/or clinically.

To explain the very rapid development of phage-resistance in the patient suffering from the *Acinetobacter* infection, both *in vivo* and *in vitro* [20, 28], we required that a fairly large fraction (*>*1%) of the pretreatment bacteria were resistant to the first cocktail (see Fig. 5). This is high compared to other estimates for the frequency of phage-resistant bacteria in bacterial populations [12, 26]. One potential explanation for this difference would be that *Acinetobacter baumannii* has been involved in a long arms race with T4-like myophages, and has evolved multiple distinct defense systems [3]. Not all defense systems need not be present in every individual bacterium, but since several are definitely expected to be present in a large bacterial population, it seems best to chose phages that are equipped to overcome those [3].

Although we have tried to use realistic parameter values by (1) adopting parameter values estimated from phage experiments in mice [26], (2) improving the estimated killing rate of neutrophils by refitting the experiments of Drusano *et al*. [10] (see Fig. S.1), (3) estimating the fitness costs and pretreatment levels of resistance by fitting time courses from *in vitro* experiments [28] (see Fig. S.5), (4) estimating the clearance rate of phages from human data [28] (see Fig. S.6), and (5) bounding the maximum replication rate of phages to *ρ* = 9.21 h^−1^, several parameter estimates remain uncertain. We also tried to develop a novel model of phage therapy in humans by (1) adopting an appropriate model developed for describing phage experiments in mice [26], (2) allowing for several phages groups and many bacterial strains being sensitive, partially resistant, or fully resistant, (3) improving the neutrophil model by making it dependent on a neutrophil source from the bone marrow, but nonetheless the model remains a caricature of the complex reality of a bacterial infection distributed over various organs. It therefore remains unclear whether or not our description of the phage therapy of the patient in Schooley *et al*. [28] is realistic. We prefer to see these simulations as examples of mechanisms that are most likely to have played a role, and we have used these results as an inspiration for further improvement of future phage therapies.

To make this model useful for interpreting future phage therapies we need better estimates of several of its crucial parameters, which requires detailed measurements during a phage therapy. For instance, knowing the bacterial resistance mechanisms for each of the cocktails [20], was essential for establishing the actual number of phage groups in our model. Measuring phage densities and bacterial strain densities longitudinally during treatment, would allow the model to better quantify the various processes determining the successfulness of future phage therapies, and help to determine exactly how such a phage therapy manages to clear a rampant bacterial infection.

Despite these uncertainties on the parameter values and mechanistic details of our models, our major observation that the outgrowth of phage-resistant bacterial strains is fast because they are released from competition with the dominant pool of sensitive bacteria, is so general that it does not depend on the details of our model. The hitting hard and early [13] with cocktails setting a high genetic barrier [2], is a recommendation applying to any species that rapidly evolves resistance to a treatment.

## 4 Acknowledgements

Most of this work was performed at the Santa Fe Institute and at the Los Alamos National Laboratory during several summer visits of RJDB to ASP. This work was supported in part by Los Alamos National Laboratory LDRD grant 20200695E (ASP).

## S.1 Supplemental figures

**Figure S.1:**
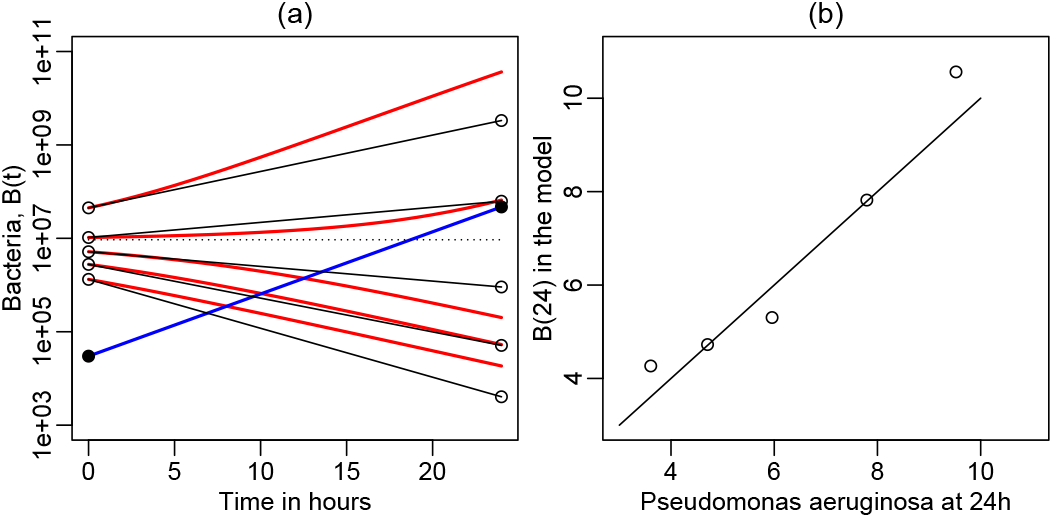
Fitting the bacterial growth data shown in Fig. 1 of Drusano *et al*. [10]. In Panel (a) the blue line depicts an example of bacterial growth in neutropenic mice (the bullets depict the observed concentrations at *t* = 0 and *t* = 24h). Assuming exponential growth this suggests a growth rate of *r* = 0.306 h^−1^ (see the main text). The black solid lines in Panel (a) depict bacterial growth curves starting at 5 different initial densities (the black circles depict the observations). Using these initial densities, the final densities were fitted with the model of Eq. (20) using *κ* and *h*_*k*_ as free parameters, i.e., for *K* = 8.48 × 10^11^ CFU/g and *r* = 0.306 h^−1^, we estimate *κ* = 0.5 and *h*_*k*_ = 1.5 × 10^7^ CFU/g. These predictions are shown as red lines in Panel (a) and as symbols in Panel (b). The horizontal dotted line in Panel (a) depicts the CBC of Eq. (20), i.e., *h*_*k*_(*κ/r* − 1), for these estimated parameters (which indeed separates densities that expand from those that contract).

**Figure S.2:**
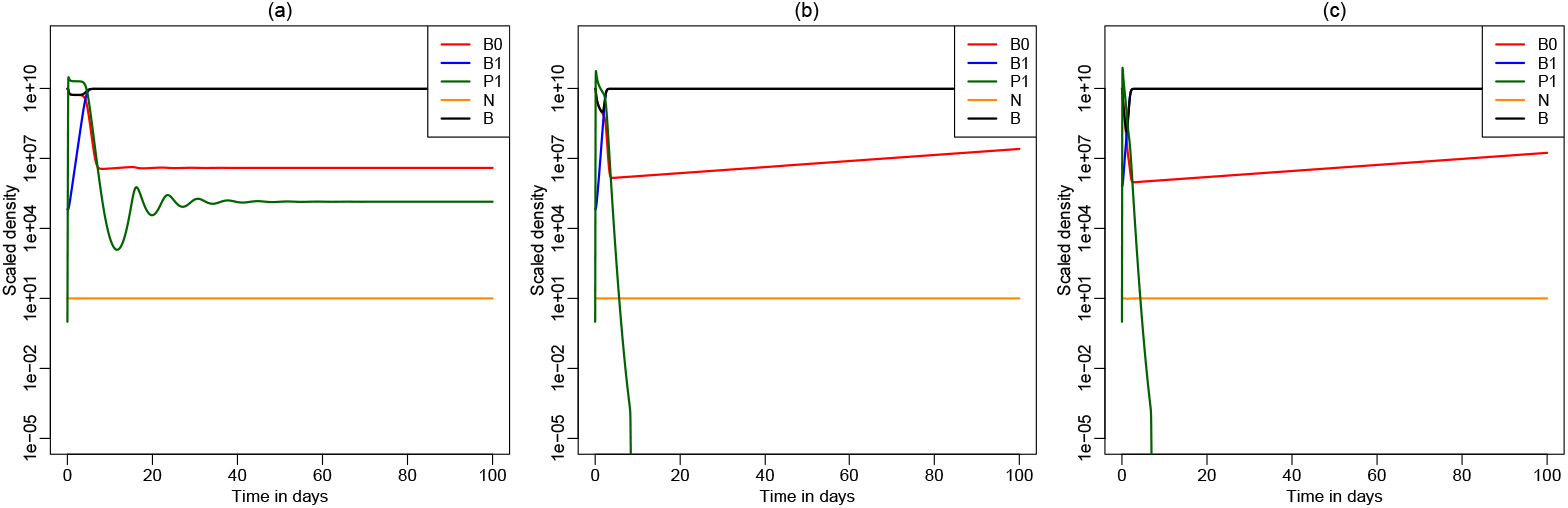
A continuation of Panels (b), (d) and (f) from Fig. 3. Panel (a) reveals that the phages recover following their initial decline, and that the system approaches a steady state with sensitive bacteria (*B*_0_), resistant bacteria (*B*_1_), and phages (*P*_1_) after a dampened oscillation that largely involves the phages and the sensitive bacteria. Panels (b) and (c) reveal that the phases go extinct during their initial decline (i.e., their densities drop below the threshold *θ* corresponding to about one phage per 10 kg of tissue).

**Figure S.3:**
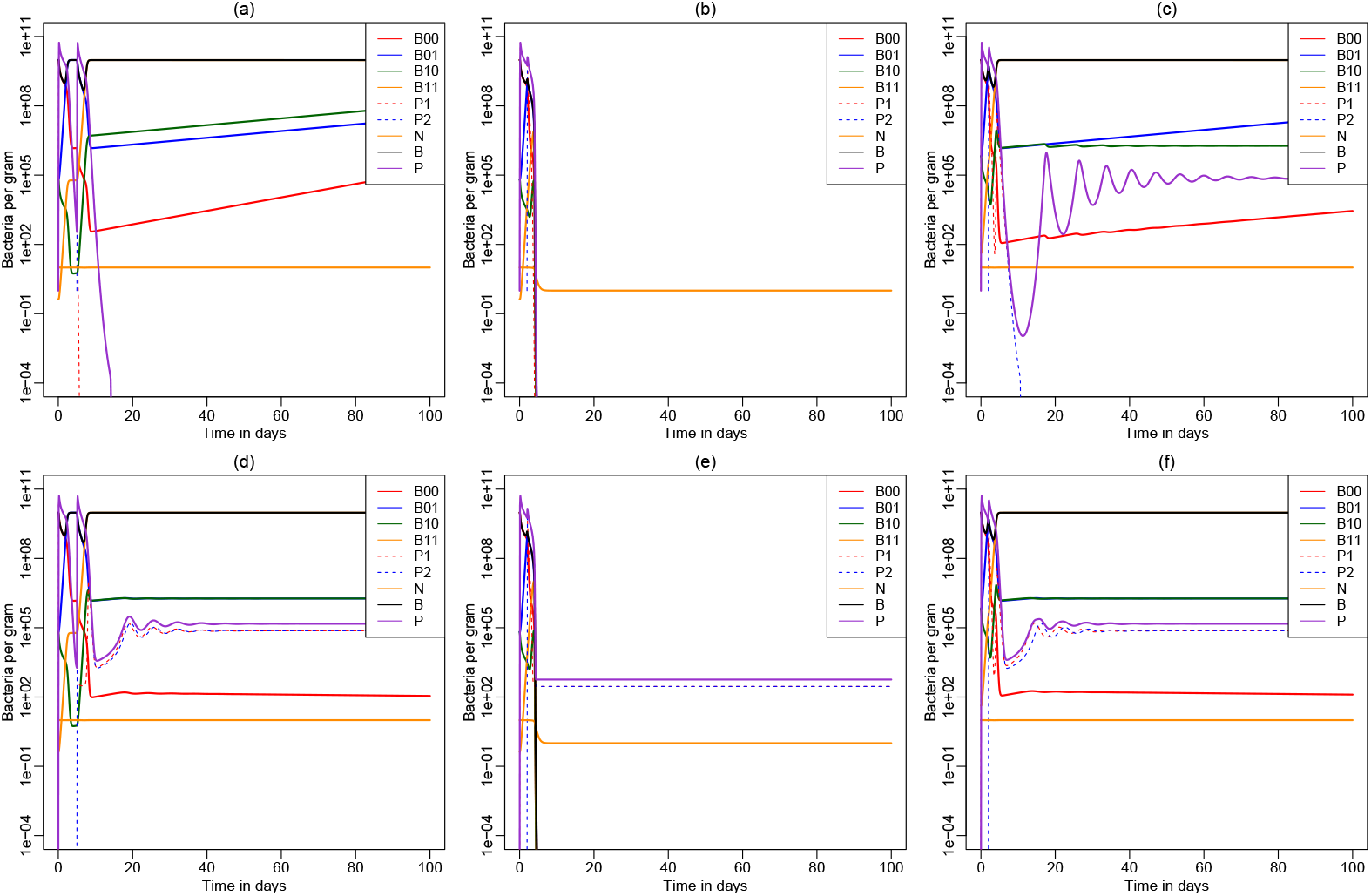
A continuation of the simulations depicted in Fig. 4. The phages go extinct in Panels (a) and (b) and persist in Panels (d – f). Phages go extinct in Panel (a) because their density drops below the extinction threshold, *θ*, when the fully resistant bacteria take over. They go extinct in Panel (b) because the bacteria are cleared. In Panel (e) where the bacterial infection is also cleared, the phages persist at a steady state determined by the ratio of infusion and loss (i.e.,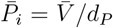). In Panels (c), (d) and (f), where the resistant bacteria breach the upper CBC, the phages largely persist by infecting a minor population of sensitive bacteria (this steady state is again approached via a dampened oscillation).

## S.2 Supplemental text

### S.2.1 Derivation of the model

To derive the infection term mechanistically we separate the time scale of phages binding bacteria from the time scale governing the cellular dynamics. First, consider a group of phages, *P*_*i*_ with *i* = 1, 2, …, *n*, that are using the same type of receptor molecule to enter and infect the bacteria. It is important to sub-divide the phages into such groups, because there will be competition amongst phages binding the same type of receptor, and if the bacteria down-regulate this receptor as a phage-resistance mechanism, it will equally affect all phages in this particular group. Next consider one bacterium that is expressing a total of *T* copies of this particular receptor molecule. The rate at which the cell will become infected by one of the phages is determined by the number of phages bound to a receptor, and becoming internalized. Using *R* for the number of unbound receptors, *a*_*i*_ for the binding (or absorption) rate of phage *i*, and *e*_*i*_ for the entry rate of phage *i*, we consider the scheme

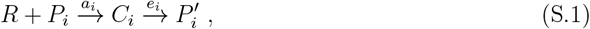

where *C*_*i*_ with *i* = 1, 2, …, *n*, defines the number of receptors bound by phages of type *i*, and 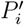 defines the corresponding number of internalized phages (see the cartoon in Fig. S.4a). The infection rate, *F*, of the bacterium is assumed to be proportional to the number of internalized phages, i.e., 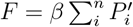.

Defining the number of unbound receptors by the conservation equation 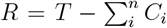 we write for the receptors bound by phages in group *i*

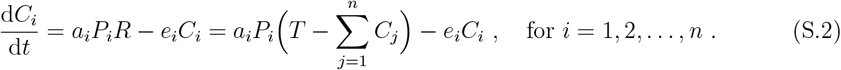

Making the QSSA d*C*_*i*_*/*d*t* = 0 for all the *n* phages binding this receptor, we arrive at

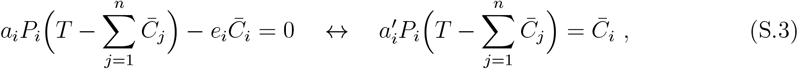

where 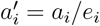. Summing these expressions over the phage groups yields

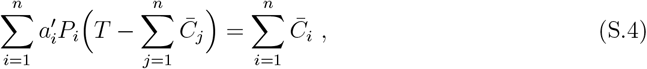

or

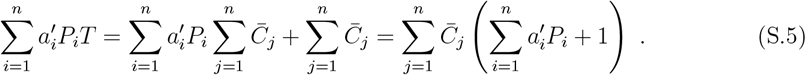

Hence

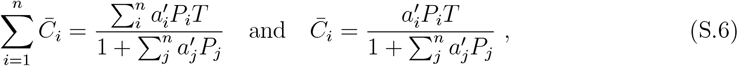

where 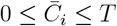. For the internalized phages we write 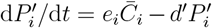, where *d*^′^ is their intracellular decay rate. Also making a QSSA 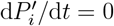 we finally arrive at

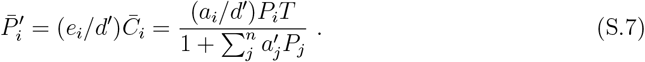

(For a single type of phage this can be written as the saturated infection rate of Eq. (1), i.e., as *βP/*(1 + *P/h*_*P*_), where *β* = *a*_1_*T/d*^′^ and *h*_*P*_ = *e*_1_*/a*_1_).

Since 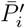 provides the number of internalized phages of type *i*, we still need to sum over all *n* phage types to arrive at the infection rate imposed by all phages binding this receptor, i.e.,

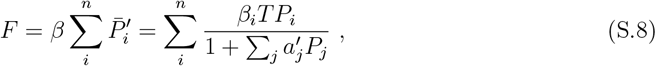

where *β* is an infection rate of the bacterium, *F* is its force of infection, and *β*_*i*_ = *βa*_*i*_*/d*^′^ combines the infection rate, with the binding rate, the entry rate, and the decay rate, and 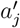 remains the ratio of the binding rate over the entry rate of phage *j* (see Fig. S.4a). Since different phage groups may bind different receptors (see Fig. S.4b), each expressed with *T*_*k*_ copies, the infection rate of Eq. (S.8) still needs to be summed over all different groups of phages, to obtain the overall infection rate, i.e.,

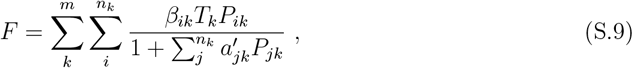

where *m* is the number of phage groups, *n*_*k*_ is the number of different phages in group *k*, and *P*_*ik*_ is density of phage *i* in group *k*, with corresponding rates *β*_*ik*_ and 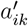.

**Figure S.4:**
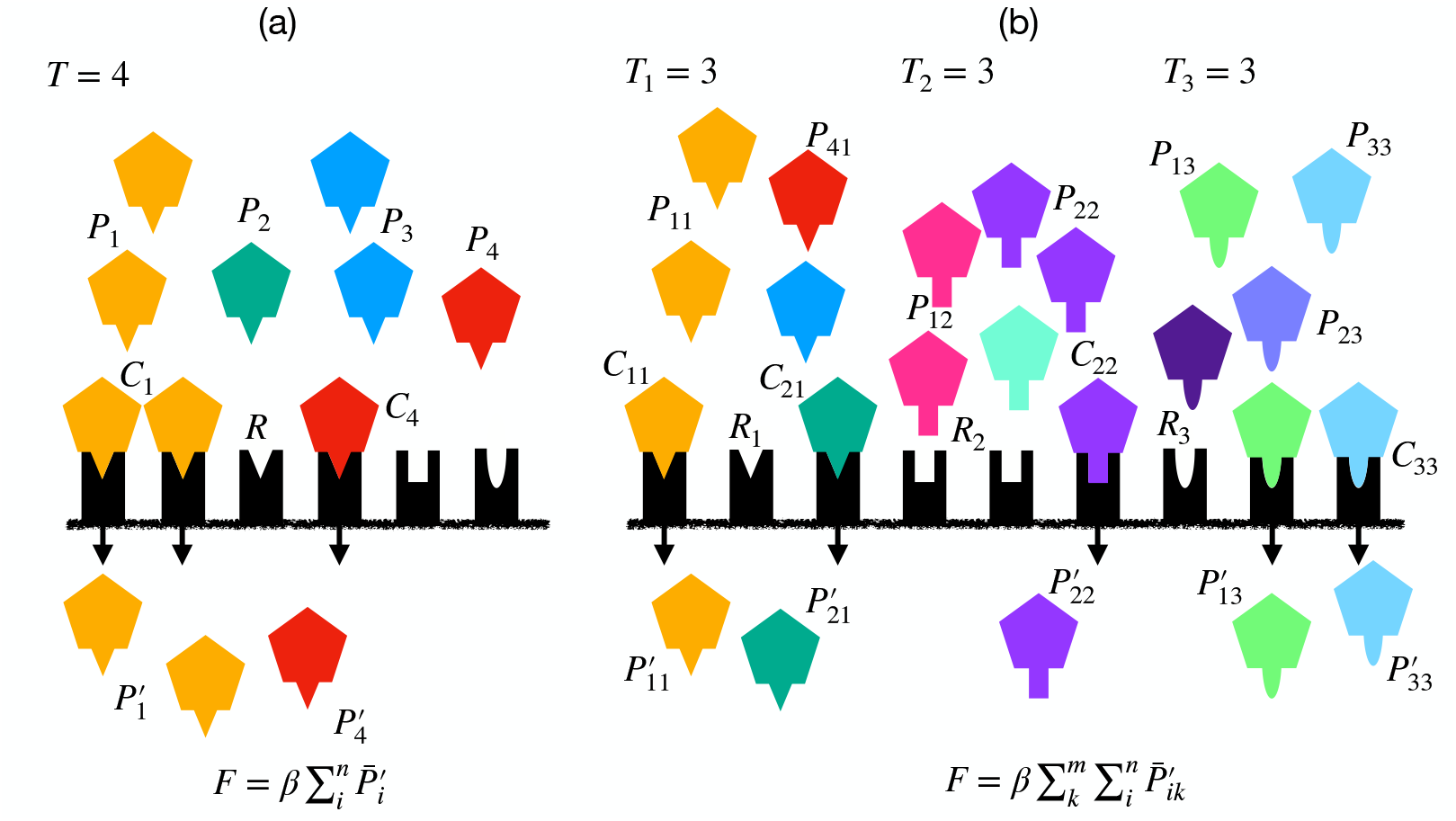
A cartoon explaining the model of Eq. (S.8) (Panel a) and Eq. (S.9) (Panel b). The heavy horizontal line depicts the bacterial cell wall. The black boxes with different indentations are receptors having different binding sites (triangular, rectangular or oval). Panel (a) depicts one group of phages, *P*_*i*_, *i* = 1, …, 4 (where the type, *i*, is reflected by the color), all interacting with the same triangular ligand, that can bind receptors with a triangular indentation. We depict a total of *T* = 4 receptors carrying a triangular binding site. Bound receptors are indicated as a complexes, *C*_*i*_, free receptors for this group as *R*, and internalized phases are called 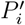. The force of infection, *F*, is proportional to the total number of internalized phages (see Eq. (S.8)). Panel (b) extends this into three phage groups, with a total of *T*_*i*_ = 3 receptors per group. Free phages *P*_*ik*_ only bind receptors of group *k* (see the complementary shapes), forming complexes *C*_*ik*_, that are internalized to become 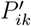 phages that all contribute to the rate of infection (see Eq. (S.9)).

Because we typically have no information about the receptor molecules the phages used in therapies bind, and since Eq. (S.9) is complicated and requires two different (typically unknown) parameters for each phage, we decided to simplify the infection term by considering all phages within one group as a single phage, having an infection rate, *β*_*i*_, and saturation constant, *e*_*i*_*/a*_*i*_. Summing over the different groups, each containing a single “super phage” the infection term becomes

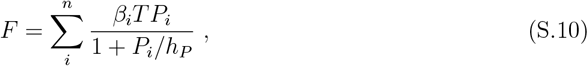

where *n* now stands for the number of groups, and we have assumed that all groups have the same saturation constant *h*_*P*_ = *e*_*i*_*/a*_*i*_, and all receptors have the same copy number, *T* = *T*_*i*_. Thus, although the infection rate of each phage group approaches a maximum *β*_*i*_*h*_*P*_, adding additional phage groups allows the total infection rate to go up to an expected value 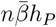, where 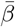 is the average infection rate.

Finally, to prevent the infection terms to become unrealistically fast when large numbers of bacteria are each expressing *T* receptors, we also allow for saturation in the bacterial densities by rewriting Eq. (S.10) into

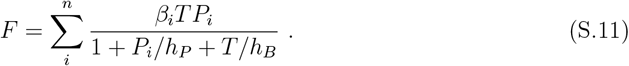

This double-saturation function can be derived mechanically for systems where a single type of phages interact with a single type of bacteria [4], as in Eq. (9), but for the high-dimensional system many types of phages and bacteria that we consider here this function remains somewhat phenomenological [7]. The main advantage of the double saturation is that the infection term approaches *h*_*B*_*β*_*i*_*P*_*i*_ when *T* → ∞, and *h*_*P*_ *β*_*i*_*T* when *P*_*i*_ → ∞.

### S.2.2 Fitting the sensitivity assays

The activity of the bacteriophage cocktails was tested by growing isolates of *A. baumannii* sampled from the patient, in the presence and absence of the phages *in vitro* (see Figure 3 in Schooley *et al*. [28]. Assuming that infected bacteria hardly replicate and that resistant bacteria are not infected, the relative growth rate observed in these experiments should reflect the relative fitness of the resistant strains. The density of the bacteria was measured over time for 20 h in “relative respirational units”, and assuming that this measure is proportional to the bacterial density, we fit a simple model for bacterial growth to these data. Finally, making the simplifying assumption that the bacterial growth rate depends linearly on the nutrient concentration, we write

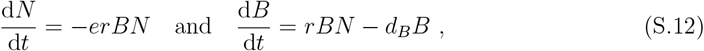

where *B* is the bacterial density, 0 *< N <* 1 the scaled nutrient concentration, *e* is the amount of resource required per bacterium, *r* the maximum replication rate, and *d*_*B*_ the death rate. Because the observed respiration initially varied around 50 units even in the absence of bacterial growth, we subtracted the minimum of the respiration observed in each experiment, and fitted *B*(*t*) to the background-corrected data (see Fig. S.5). We fit the data in each experiment simultaneously allowing only the replication rate, *r*, and the initial condition, *B*(0), to vary between the bacteria with and without all the phages (see the red and black lines in Fig. S.5, respectively). Differences in the replication rate should reflect fitness costs, and differences in the initial condition should reflect initial densities of phage-resistant bacterial strains present within the isolates.

When we fit the model of Eq. (S.12) to the data of bacteria growing in the absence of phages we obtain an average replication rate of the various strains weighted by their frequencies (the match between the black symbols and the black lines in Fig. S.5 show that the model describes the data well). In the presence of phages (red symbols in Fig. S.5), there is initially hardly any expansion of the bacteria, which provides support for our assumption that in these cultures sensitive bacteria are infected so rapidly that they fail to expand, and hence that the estimated replication rate should reflect the growth rate of strains that are resistant to the phages. The model also describes these data well; see Table S.1 for the parameter estimates.

The first data set of Schooley *et al*. [28] involved growing bacteria isolated from the patient before the onset of phage therapy with or without the phages comprising the first cocktail *ϕ*PC (see Fig. 3a in their paper, and Fig. S.5a of this paper). The model fits the data with and without *ϕ*PC phages quite well (see Fig. S.5a), and provides reasonable parameter estimates (Table S.1), i.e., a doubling time of about an hour, 0.43 *< r <* 0.58 h^−1^, and a negligible death rate, *d*_*B*_ = 0.042 h^−1^. The bacteria resistant to the full cocktail have an estimated replication rate, *r* = 0.43 h^−1^, which is about 74% of that of the average bacterial population having *r* = 0.58 h^−1^ (i.e., in Table S.1 we write *ρ* = 0.43*/*0.58 = 0.74). Since this is the relative growth with respect to the *B*_00_ ‘wild type’ having a fitness *ϕ*_1_ = 1, we set *ϕ*_2_ = 0.74 for the *B*_01_ strain. The initial density of the resistant phages is *B*(0) = 1.01, which is about 21% of that of the total *B*(0) = 4.89.

**Figure S.5:**
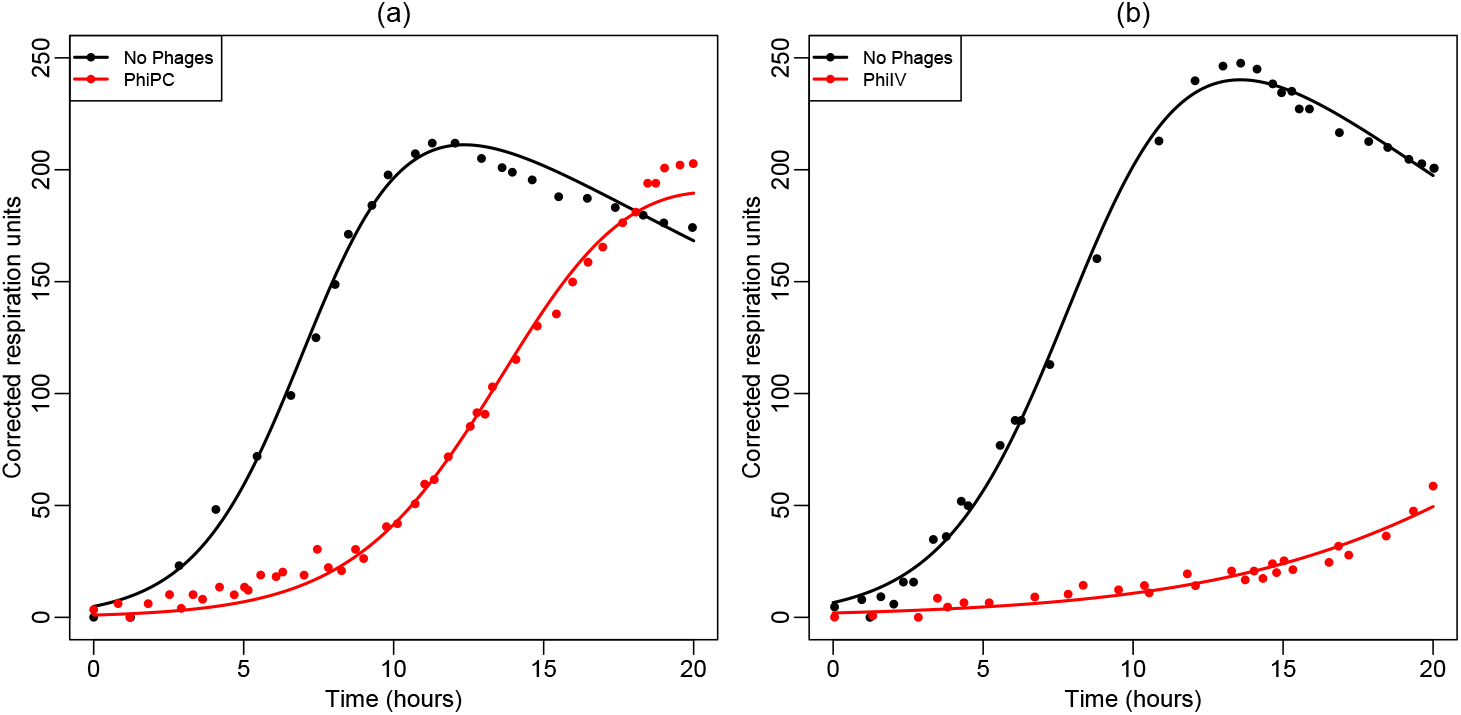
Activity of the bacteriophage cocktails on bacterial isolates taken before treatment (Panel a) or after six days of treatment (Panel b). Symbols depict the data copied from Fig. 3 in Schooley *et al*. [28] and lines depict the best fits of Eq. (S.12) to this data. The parameter estimates are provided in Table S.1. Fitting was performed by using the Levenberg-Marquardt algorithm implemented in FME [31].

The second data set of Schooley *et al*. [28] is similar except that it uses bacteria isolated from the patient at day six and the second “Navy” cocktail, *ϕ*IV (see Fig. 3f in their paper). These data are reproduced in Fig. S.5b for the bacteria only (black bullets), and for bacteria in the presence of the full *ϕ*IV cocktail (red). The model also fits these two data sets reasonably well (see Fig. S.5b), and provides similar parameter estimates (Table S.1), i.e., a doubling time of about an hour in the absence of phages *r* 0.51 h^−1^, and a negligible death rate, *d*_*B*_ = 0.051 h^−1^. In the presence of phages we estimate a replication rate of *r* = 0.22 h^−1^, or a relative growth rate of *ϕ*_2_ = 0.22*/*0.51 = 0.44 h^−1^. Since this is the relative growth with respect to the *B*_01_ strain having a fitness *ϕ*_2_ = 0.74, we set *ϕ*_4_ = 0.74 × 0.44 = 0.33 for the *B*_11_ strain. At day six the density of the resistant phages is *B*(0) = 1.95, which is about 30% of that of the total *B*(0) = 6.56.

### S.2.3 Fitting the bacteriophage pharmacokinetics

The decay rate of phages in human serum was measured after the administration of 5 × 10^9^ PFU of bacteriophage via intravenous injection (see Fig. 4 in Schooley *et al*. [28]). We depict this data in Fig. S.6 by green circles, and fit them in two ways with the following model

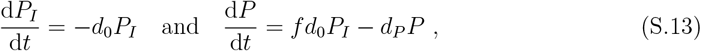

where we track the concentration of the phages in the inoculum, *P*_*I*_, and phages in the serum, *P*. Phages are lost from the inoculum at rate *d*_0_, a fraction *f* appears in the serum, where they are lost at rate *d*_*P*_. Setting *P*_*I*_(0) = 5 × 10^9^ and *P* (0) = 0 we fitted *P* (*t*) as predicted by Eq. (S.13) to the observed plasma levels, and estimated *d*_0_ = 16 h^−1^, *d*_*P*_ = 4 h^−1^, and *f* = 5 × 10^−6^. This best fit is depicted as the red line in Fig. S.6. We also fitted the same data with simple exponential decay by setting *I*(0) = 0, and estimated *P* (0) = 2.4 × 10^4^ and *d*_*P*_ = 3.5 h^−1^, which is shown as the blue line in Fig. S.6. Since these two estimates of *d*_*P*_ are similar, and the fit of the exponential decay model is somewhat better, we use the estimate based upon the simplest model, *d*_*P*_ = 3.5 h^−1^, throughout (see Table S.2).

**Table S.1:** Fitting the respiration data of Schooley *et al*. [28]. The values between the brackets denote standard errors computed from the Hessian matrix by the FME package [31]. Common parameters were similar for the pretreatment and the day-six experiments, i.e., *d*_*B*_ = 0.042 and *e* = 0.0036, and *d*_*B*_ = 0.051 and *e* = 0.0029, respectively.

| Parameter | Pretreatment |  | Day six |  |
| --- | --- | --- | --- | --- |
| | No phages | Cocktail ΦPC | No phages | Cocktail ΦIV |
| $r$ ($\text{h}^{-1}$) | 0.58 ($\pm 0.03$) | 0.43 ($\pm 0.01$) | 0.51 ($\pm 0.01$) | 0.22 ($\pm 0.02$) |
| $B(0)$ | 4.49 ($\pm 1.05$) | 1.01 ($\pm 0.15$) | 6.56 ($\pm 0.68$) | 1.95 ($\pm 0.51$) |
| Growth rate $\rho$ | 0.43/0.58 = 0.74 | | 0.22/0.51 = 0.44 | |
| Fitness $\phi_i$ | $\phi_2 = 0.74$ | | $\phi_4 = 0.74 \times 0.44 = 0.33$ | |
| Frequency | 1.01/4.49 = 0.21 |  | 1.95/6.56 = 0.30 |  |

**Figure S.6:**
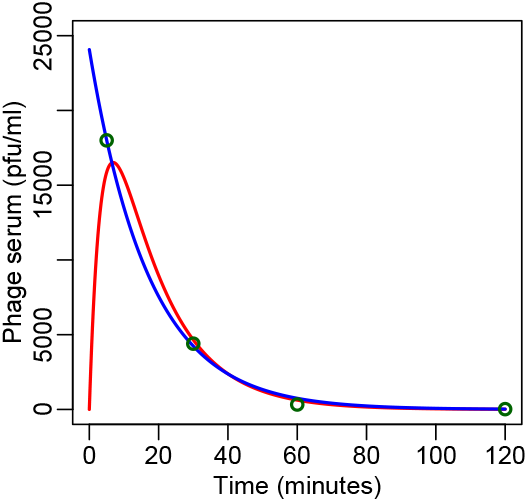
The phage titers measured by Schooley *et al*. [28]. The symbols depict the serum phage concentration data copied from Fig. 4 in Schooley *et al*. [28], the red line the results of fitting of this data with Eq. (S.13), and the blue line the result of fitting with an exponential decay model.

**Table S.2:** Parameter values of the human model.

| Name | Value | Dimension | Meaning | Reference |
| --- | --- | --- | --- | --- |
| $r$ | 0.31 | $\text{h}^{-1}$ | Bacterial replication rate | Table 1 |
| $d_B$ | 0.01 | $\text{h}^{-1}$ | Bacterial death rate | Guessed |
| $b$ | 100 | — | Burst size | [26] |
| $K$ | $10^{10}$ | CFU/g | Carrying capacity | [26] |
| $d_I$ | 2 | $\text{h}^{-1}$ | lysis rate bacteria | [22, 35] |
| $d_P$ | 3.5 | $\text{h}^{-1}$ | Clearance rate phages | Fig. S.6 |
| $k$ | $1.5 \times 10^{-7}$ | $\text{h}^{-1}$ | Killing rate by neutrophils | Table 1 & [26] |
| $h_N$ | $10^7$ | CFU/g | Saturation neutrophil increase | [26] |
| $h_k$ | $2.86 \times 10^6$ | CFU/g | Saturation neutrophil killing | Table 1 |
| $h_P$ | $1.5 \times 10^7$ | PFU/g | Saturation infection rate | [26] |
| $h_B$ | $3.56 \times 10^7$ | CFU/g | Saturation infection rate | Eq. (17) |
| $d_N$ | 0.0833 | $\text{h}^{-1}$ | Time scale of neutrophils ($d_N = 0.5 \text{ d}^{-1}$) | [1, 33] |
| $\sigma$ | $2.25 \times 10^5$ | cells $\text{h}^{-1}$ | $\sigma = N(0)/d_N$ | [26] |
| $\alpha$ | 9 | — | Defines a 10-fold increase of neutrophils | [25, 26] |
| $cA_i$ | 0 | $\text{h}^{-1}$ | Clearance rate by antibody $i$ | — |
| $a$ | 0 | $\text{h}^{-1}$ | Absorption rate | — |
| $\beta_{\text{crit}}$ | $2 \times 10^{-8}$ | g/(PFU h) | Minimal infection rate | Computed |
| $\bar{\phi}$ | 0.33 – 1 | — | Relative fitnesses | Table S.1 |
| $\mu$ | $2.85 \times 10^{-8}$ | — | Mutation rate in Fig. 3 | [26] |
| $\mu$ | 0.0005 – 0.05 | — | Mutation rates in Fig. 5 & 6 | Table S.1 |
| $\bar{V}$ | $10^3$ | $\text{h}^{-1}$ | Phage infusion rate | Guessed |

## Footnotes

1 To allow for clearance, one could also consider *I*(1) = 10^3^, and have a lower bound *ρ* = 6.91 h^−1^.

2 Note that we do not adjust *h*_*B*_ when 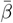 changes, and that phages with an infection rate exceeding *β*_crit_, will therefore initially replicate faster than *ρ* = 9.21 h^−1^. This does not affect our result that increasing the infection rate hardly matters, because these initial phage replication rates allow for even better control.

